# The Disease-Associated Proteins *Drosophila* Nab2 and Ataxin-2 Interact with Shared RNAs and Coregulate Neuronal Morphology

**DOI:** 10.1101/2021.03.01.433469

**Authors:** J. Christopher Rounds, Edwin B. Corgiat, Changtian Ye, Joseph A. Behnke, Seth M. Kelly, Anita H. Corbett, Kenneth H. Moberg

## Abstract

*Nab2* encodes a conserved polyadenosine RNA-binding protein (RBP) with broad roles in post-transcriptional regulation, including in poly(A) RNA export, poly(A) tail length control, transcription termination, and mRNA splicing. Mutation of the *Nab2* human ortholog *ZC3H14* gives rise to an autosomal recessive intellectual disability, but understanding of Nab2/ZC3H14 function in metazoan nervous systems is limited, in part because no comprehensive identification of metazoan Nab2/ZC3H14-associated RNA transcripts has yet been conducted. Moreover, many Nab2/ZC3H14 functional protein partnerships likely remain unidentified. Here we present evidence that *Drosophila melanogaster* Nab2 interacts with the RBP Ataxin-2 (Atx2), a neuronal translational regulator, and implicate these proteins in coordinate regulation of neuronal morphology and adult viability. We then present the first high-throughput identifications of Nab2- and Atx2-associated RNAs in *Drosophila* brain neurons using an RNA immunoprecipitation-sequencing (RIP-Seq) approach. Critically, the RNA interactomes of each RBP overlap, and Nab2 exhibits high specificity in its RNA associations in neurons *in vivo*, associating with a small fraction of all polyadenylated RNAs. The identities of shared associated transcripts (e.g. *drk*, *me31B*, *stai*) and of transcripts specific to Nab2 or Atx2 (e.g. *Arpc2*, *tea*, respectively) promise insight into neuronal functions of and interactions between each RBP. Significantly, Nab2-associated RNAs are overrepresented for internal A-rich motifs, suggesting these sequences may partially mediate Nab2 target selection. Taken together, these data demonstrate that Nab2 opposingly regulates neuronal morphology and shares associated neuronal RNAs with Atx2, and that *Drosophila* Nab2 associates with a more specific subset of polyadenylated mRNAs than its polyadenosine affinity alone may suggest.

## Introduction

Intellectual disability refers to a broad group of neurodevelopmental disorders affecting approximately 1% of the world population (Maulik *et al*. 2011) and defined by significant limitations in intellectual functioning and adaptive behavior (Tassé *et al*. 2016; Vissers *et al*. 2016). Intellectual disabilities are etiologically diverse and in some cases genetically complex, yet many exhibit overlapping molecular dysfunctions in a comparatively limited set of fundamental neurodevelopmental pathways (reviewed in Chelly *et al*. 2006; van Bokhoven 2011; and Verma *et al*. 2019). Thus, monogenic intellectual disabilities represent experimentally tractable avenues for understanding both these disorders more broadly and neurodevelopment in general (Najmabadi *et al*. 2011; Agha *et al*. 2014). One set of such informative monogenic intellectual disabilities is caused by mutations affecting genes encoding RNA-binding proteins (RBPs) (reviewed in Bardoni *et al*. 2012) such as *ZC3H14* (z*inc finger CCCH-type containing 14*). Specifically, loss-of-function mutations in *ZC3H14*, which encodes a ubiquitously expressed polyadenosine RBP, cause a non-syndromic form of autosomal recessive intellectual disability (Pak *et al*. 2011; Al-Nabhani *et al*. 2018). However, the molecular functions and developmental roles of human ZC3H14 are largely unknown; defining these functions and roles provides an opportunity to better understand intellectual disability and human neurodevelopment.

*Drosophila melanogaster* has proven a powerful model system to understand the molecular functions of proteins encoded by many intellectual disability genes (Inlow and Restifo 2004; Oortveld *et al*. 2013), and ZC3H14 is no exception—its functions have begun to be dissected in part through study of its *Drosophila* ortholog Nab2 (Pak *et al*. 2011; Kelly *et al*. 2014). *Drosophila* Nab2, like ZC3H14, is a polyadenosine RNA-binding protein that induces neurological defects when its expression is altered; deletion or overexpression of *Nab2* causes neuronal morphological defects in the eye, axon projection defects in the developing brain, and memory impairments (Pak *et al*. 2011; Kelly *et al*. 2016; Bienkowski *et al*. 2017; Corgiat *et al*. 2020). The function of Nab2 is particularly important in *Drosophila* neurons, as pan-neuronal expression of Nab2 or an isoform of human ZC3H14 is sufficient to rescue the severe limitation in adult viability and locomotor defects caused by zygotic Nab2 deficiency (Pak *et al*. 2011; Kelly *et al*. 2014). Crucially, Nab2 physically and functionally interacts with Fmr1, the *Drosophila* homolog of the Fragile X Syndrome RBP FMRP (Verkerk *et al*. 1991; Ashley *et al*. 1993; Wan *et al*. 2000), to support axonal morphology and olfactory memory (Bienkowski *et al*. 2017). Previous data suggest functions of *Drosophila* Nab2 in poly(A) tail length control, translational regulation, and mRNA splicing, but mechanistic demonstrations of its molecular function on individual, endogenous transcripts have yet to emerge (Pak *et al*. 2011; Kelly *et al*. 2014; Bienkowski *et al*. 2017; Jalloh *et al*. 2020). Such demonstrations have been prevented in large part because very few *Drosophila* Nab2-associated RNAs have been identified (Bienkowski *et al*. 2017; Jalloh *et al*. 2020), and a comprehensive accounting of Nab2-associated RNAs has yet to be conducted.

While the precise molecular function of *Drosophila* Nab2 on its associated transcripts is unknown, informed hypotheses may be drawn by synthesizing research on *Drosophila* Nab2 and orthologs murine ZC3H14, human ZC3H14, and *S. cerevisiae* Nab2, the most well-studied Nab2/ZC3H14 ortholog (reviewed in Fasken *et al*. 2019). In *S. cerevisiae*, Nab2 functions pervasively across many RNAs in transcript stability and transcription termination, and it likely acts similarly broadly in poly(A) tail length control and poly(A) RNA export (Schmid *et al*. 2015; Fasken *et al*. 2019; Alpert *et al*. 2020). Mutation of *S. cerevisiae* Nab2 induces dramatic increases in bulk poly(A) tail length and disrupts bulk poly(A) export from the nucleus (Green *et al*. 2002; Kelly *et al*. 2010). Consistent with its pervasive effects on many transcripts, *S. cerevisiae* Nab2 exhibits a broad binding target profile and is essential for cellular viability (Anderson *et al*. 1993; Tuck and Tollervey 2013). By contrast, mutant analyses of metazoan Nab2/ZC3H14 imply increased RNA target specificity for these proteins. Unlike Nab2 in *S. cerevisiae*, full-length ZC3H14 in mice and humans is not essential for viability—instead, loss of ZC3H14 decreases viability in mice and causes neurological or neurodevelopmental defects in both organisms (Pak *et al*. 2011; Rha *et al*. 2017b; Al-Nabhani *et al*. 2018). Bulk poly(A) tail lengths increase upon Nab2 loss in *Drosophila* or full-length ZC3H14 loss in mice *in vivo*, but this increase is not observed across all mouse tissues or all individual *Drosophila* mRNAs tested, and it is less pronounced than the effects observed in *S. cerevisiae* (Kelly *et al*. 2010; Bienkowski *et al*. 2017; Rha *et al*. 2017b). Moreover, in *Drosophila* and mouse cells, respectively, a pervasive nuclear poly(A) export defect is not observed upon Nab2 loss or ZC3H14 knockdown (Farny *et al*. 2008; Pak *et al*. 2011; Kelly *et al*. 2014). *Drosophila* Nab2 is required for proper splicing of individual introns and exons, but in a small, specific set of transcripts, including *Sex lethal* (Jalloh *et al*. 2020). Taken together, these data are consistent with a focused role for *Drosophila* Nab2 in regulating poly(A) tail length, splicing, stability, and nuclear export crucial for certain transcripts, cell types, and developmental contexts (Bienkowski *et al*. 2017; Rha *et al*. 2017b; Jalloh *et al*. 2020). Crucially however, the theme of *Drosophila* Nab2 RNA target specificity implied by these data has not been tested and remains an important open question, especially as the polyadenosine affinity of *Drosophila* Nab2 (Pak *et al*. 2011) makes it theoretically capable of associating with all polyadenylated transcripts through their poly(A) tails. Thus, a comprehensive identification of *Drosophila* Nab2-associated RNAs is necessary to determine the potential scope of Nab2 function and provide sets of transcripts on which the molecular consequences of Nab2-RNA association may be systematically evaluated. In the present study, in response we define the first neuronal RNA interactome for Nab2.

Contextualizing Nab2-RNA associations requires further definition of the molecular pathways and proteins, particularly other RBPs, that Nab2 interacts with or regulates. Notably, the *Nab2* modifier eye screen that initially linked Nab2 and Fmr1 (Bienkowski *et al*. 2017) also recovered an allele of *Ataxin-2* (*Atx2*), which encodes a conserved RBP and regulatory partner of Fmr1 in *Drosophila* neurons (Sudhakaran *et al*. 2014; Jiménez-López and Guzmán 2014). The shared connection of Nab2 and Atx2 with Fmr1 raised the possibility of cooperation or competition between these two proteins. Underscoring the value of this approach, Atx2 is a protein of particular importance for human health and neuronal function. Expansion of a polyglutamine tract within ATXN2, the human Atx2 ortholog, gives rise to the autosomal dominant neurodegenerative disease spinocerebellar ataxia type 2 (SCA2) (Imbert *et al*. 1996; Pulst *et al*. 1996; Sanpei *et al*. 1996). Expansions of the same tract are also associated with parkinsonism and amyotrophic lateral sclerosis (ALS) (Gwinn-Hardy *et al*. 2000; Elden *et al*. 2010; Park *et al*. 2015). Functionally, *Atx2* encodes a conserved RNA-binding protein that regulates protein translation, mRNA stability, and mRNP granule formation and plays roles in memory, cellular metabolism, and circadian rhythms (reviewed in Ostrowski *et al*. 2017; Lee *et al*. 2018). Among the most well-studied molecular roles of Atx2 are its contributions to regulation of mRNA translation in the cytoplasm. Specifically, Atx2 suppresses the translation of some target RNAs through RNP granule formation and interactions with the RNAi machinery (McCann *et al*. 2011; Sudhakaran *et al*. 2014; Bakthavachalu *et al*. 2018) and supports the translation of other targets by promoting RNA circularization (Lim and Allada 2013; Zhang *et al*. 2013; Lee *et al*. 2017). Intriguingly Atx2, like Nab2, contributes to poly(A) tail length control in *S. cerevisiae*—the yeast Atx2 ortholog Pbp1 promotes poly(A) tail length, likely by inhibiting the activity of poly(A) nuclease (PAN) (Mangus *et al*. 1998, 2004). The shared connections of Nab2 and Atx2 to Fmr1, neuronal translation, and poly(A) tail length control emphasize the potential for and need to test whether these RBPs functionally interact beyond the initial eye screen link.

Here, after expanding the genetic link previously identified between Nab2 and Atx2 in our modifier screen, we used genetic and molecular approaches to probe the functional connections between these two RBPs. We show that Nab2 and Atx2 functionally interact to control neuronal morphology of the mushroom bodies (MBs), a learning and memory center of the *Drosophila* brain (Heisenberg 2003; Kahsai and Zars 2011; Yagi *et al*. 2016; Takemura *et al*. 2017). We then present the first high-throughput identification of Nab2-and Atx2-associated RNAs in *Drosophila*; in fact, such accounting has been performed for Nab2 only in *S. cerevisiae*, not in any metazoan (Guisbert *et al*. 2005; Batisse *et al*. 2009; Tuck and Tollervey 2013; Baejen *et al*. 2014). This approach demonstrates Nab2 and Atx2 associate with an overlapping set of RNA transcripts in fly brains and provides insight into the functions of each protein individually and in concert with one another. Considering these data as a whole, we propose a model in which the genetic interactions between Nab2 and Atx2 are explained by their counterbalanced regulation of shared associated RNAs. Our data represent a valuable resource for understanding the neuronal roles of Nab2 and Atx2 in *Drosophila* and, potentially, for understanding links between each RBP and human disease.

## Materials and Methods

### *Drosophila* genetics and husbandry

Genetic crosses of *Drosophila melanogaster* were raised on standard media and maintained at 25°C in humidified incubators (SRI20PF, Shel Lab) with 12-hour light-dark cycles unless otherwise specified. Cultures were often supplemented with granular yeast (Red Star Yeast) to encourage egg laying. Parental stocks were maintained at either at room temperature (RT) or 18°C to control virgin eclosion timing. Stocks used include *Nab2^ex3^* (a *Nab2* null), *Nab2^pex41^* (a P-element excision control serving as a *Nab2* wild type), and *UAS>Nab2-FLAG*, all first described in (Pak *et al*. 2011). Additional stocks used include *GMR-Gal4* (on chromosome 2), *Atx2^X1^* (an *Atx2* null, gift of N. Bonini) (Satterfield *et al*. 2002), and *UAS>Atx2-3xFLAG* (gift of R. Allada) (Lim and Allada 2013). Finally, stocks sourced from the Bloomington Drosophila Stock Center (BDSC) include: *elav>Gal4* (*elav^c155^*, BL458) (Lin and Goodman 1994), *OK107-Gal4* (BL854) (Connolly *et al*. 1996), *Df(3R)Exel6174* (BL7653) (Parks *et al*. 2004), *UAS>Nab2* (*Nab2^EP3716^*, BL17159) (Rørth *et al*. 1998; Bellen *et al*. 2004), and *Atx2^DG08112^*. The *Atx2^DG08112^* stock (Huet *et al*. 2002) was mapped as part of the Gene Disruption Project (GDP) (Bellen *et al*. 2004) and is no longer available from the BDSC; copies provided upon request.

### *Drosophila* eye imaging

*Drosophila* eyes were imaged using a Leica MC170 HD digital camera mounted on a Nikon SMZ800N stereo microscope at 8X magnification. To prepare subjects for imaging, flies were flash frozen (−80°C, 1 minute), fixed in place on a clear Slygard pad using minutien pins (26002-10, Fine Science Tools), and submerged in 70% ethanol to diffuse light and reduce glare. Subjects were illuminated with a fiber optic ring light (Dolan-Jenner) and LED illuminator (Nikon Instruments Inc.) and image acquisition was performed using the Leica Application Suite (v4.12) for Windows under the following parameters: 140 ms exposure; automatic white balance; highest available resolution; and default values for gain, saturation, gamma, and hue. Each subject was imaged at multiple focal planes (often ≥ 10), and these were subsequently combined using the *Auto-Align* and *Auto-Blend* functions in Photoshop CS5.1 Extended (Adobe) to generate final, merged images in which the entire subject is in-focus. These “focus stacking” processing steps (Patterson) combine only in-focus regions of an image series into a single, merged image.

### Immunofluorescence

For mushroom body morphology experiments, *Drosophila* brains were dissected using methods similar to those in (Williamson and Hiesinger 2010; Kelly *et al*. 2016, 2017). Briefly, using #5 Dumont fine forceps (Ted Pella, Inc.), for each dissection a *Drosophila* head was isolated in PBS (often supplemented with 0.1% Triton X-100), the proboscis was removed to provide a forceps grip point, and the remaining cuticle and trachea were peeled away from the brain within. On wet ice, dissected brains were fixed in 4% paraformaldehyde for 30 minutes and then permeabilized in 0.3% PBS-Triton (PBS-T) for 20 minutes. For both primary and secondary antibody incubations, brains were left rocking at 4°C for 1-3 nights in 0.1% PBS-T supplemented with blocking agent normal goat serum (Jackson ImmunoResearch) at a 1:20 dilution. Immunostained brains were mounted on SuperFrost Plus slides (12-550-15, Fisher Scientific) in Vectashield (H-1000, Vector Laboratories) using a cover slip “bridge” method (Kelly *et al*. 2017). Brains were imaged on a Zeiss LSM 510 confocal microscope. Exclusively female flies were dissected for practicality, given that *Nab2^ex3^* nulls were analyzed in this experiment and *Nab2^ex3^* adult viability skews towards females (Jalloh *et al*. 2020).

For Nab2-Atx2 localization experiments, whole animals were fixed in 4% paraformaldehyde, 0.008% PBS-T, shaking, for 3 hours at RT and then washed in PBS and stored at 4°C overnight. Brains were dissected in 0.008% PBS-T using similar methods as described above, permeabilized by shaking in 0.5% PBS-T overnight at 4°C, and blocked by shaking in 0.5% PBS-T, 5% NGS for 2 hours at RT. For both primary and secondary antibody/Hoechst incubations, brains were left shaking at 4°C for 2-3 nights in 0.5% PBS-T, 5% NGS. After washing with 0.5% PBS-T followed by PBS, brains were mounted in SlowFade Gold Antifade Mountant (S36936, Invitrogen), surrounded by an adhesive imaging spacer (GBL654002, Sigma-Aldrich) to prevent sample compression, and finally cover-slipped and sealed with clear nail polish. Brains were imaged on an A1R HD25 confocal microscope (Nikon) and a multi-photon FV1000 laser-scanning microscope (Olympus).

Primary antibodies and dilutions used are as follows: mouse α-Fasciclin 2 (1:50) (1D4, Developmental Studies Hybridoma Bank), rabbit α-GFP (1:400) (A11122, Invitrogen), and mouse α-FLAG (1:500) (F1804, Sigma-Aldrich). Secondary antibodies and dilutions used are as follows: goat α-mouse Cy3 (1:100) (Jackson ImmunoResearch), goat α-mouse Alexa 594 (1:400) (A11032, Invitrogen) and goat α-rabbit Alexa 488 (1:400) (A11008, Invitrogen). To fluoresce DNA and mark nuclei in localization experiments, brains were also incubated with a Hoechst 33342 stain (1:1,000) (H21492, Invitrogen) during secondary antibody incubation.

Further brain image analysis and processing, including generating maximum intensity projections and focus stacks and adjusting brightness and contrast, was performed with Photoshop CS5.1 Extended (Adobe) and Fiji (Schindelin *et al*. 2012), a distribution of ImageJ (Schneider *et al*. 2012; Rueden *et al*. 2017).

### Immunoprecipitation

This immunoprecipitation protocol was developed through optimization guided by the protocols presented in (Yang *et al*. 2005; Banerjee *et al*. 2017; Bienkowski *et al*. 2017; Morris and Corbett 2018). Nuclear Isolation Buffer (NIB; 10 mM Tris HCl pH 7.4, 10 mM NaCl, 3 mM MgCl_2_, 0.5% NP-40) and Immunoprecipitation Buffer (IP Buffer; 50 mM HEPES, 150 mM NaCl, 5 mM EDTA, 0.1% NP-40) were prepared ahead of the experiment and stored indefinitely at 4°C. Both buffers, and the glycine and PBS solutions below, were prepared primarily in 0.1% diethyl pyrocarbonate (DEPC)-treated and autoclaved ultrapure Milli-Q water to limit RNase contamination. Both NIB and IP Buffer were supplemented with an EDTA-free cOmplete protease inhibitor cocktail tablet (1 tablet/28 ml; 11873580001, Roche) and RNasin Plus RNase inhibitor (0.2%; N2615, Promega) freshly before each experiment. Additionally, before each experiment Protein G-coupled magnetic Dynabeads (10003D, Thermo Fisher) were conjugated to glycerol-free (Domanski *et al*. 2012) monoclonal α-FLAG (F3165, Sigma-Aldrich) in aliquots of 1.5 mg beads/9 µg antibody by incubation for 45 minutes at room temperature. Throughout the experiment, beads were magnetized using DynaMag-Spin magnets (e.g. 12320D, Thermo Fisher) as necessary. Exclusively female flies were used for consistency with MB experiments and for practicality, as both *elav>Nab2-FLAG* and *elav>Atx2-3xFLAG* prohibitively decreased relative male viability (data not shown), presumably due to deleterious effects in males likely driven by dosage compensation of the X-chromosome-linked *elav>Gal4* construct leading to enhanced epitope-tagged protein overexpression.

300 female *Drosophila* heads each of the genotypes *elav>Gal4* alone, *elav>Nab2-FLAG*, and *elav>Atx2-3xFLAG*, previously isolated in bulk (see *Supplemental Materials and Methods*), were fixed in 1% formaldehyde, 0.1% NP-40 in PBS for 30 minutes at 4°C. Fixation was quenched by adding glycine to a final concentration of 250 mM and rocking for 10 minutes at 4°C. Heads were washed in 0.1% NP-40 in PBS and then manually homogenized with a smooth Teflon pestle for 5 minutes in 250 µL of NIB in a size AA glass tissue grinder at 4°C (3431D70, Thomas Scientific). Homogenates were spun through 35 µm cell strainer caps into round-bottom tubes (352235, Falcon) to remove exoskeletal debris, transferred, and then centrifuged for 5 minutes at 500×g at 4°C to separate an insoluble fraction. Twenty percent of the soluble supernatant volume was isolated and defined as Input; the remaining eighty percent was used for immunoprecipitation. Both Input and IP samples were diluted to final concentrations of 0.8x IP Buffer to ensure comparable and efficient sample lysis. IP samples were transferred onto the α-FLAG-conjugated magnetic Dynabeads, and both sample types were incubated, rotating, for 10 minutes at room temperature. Next, IP sample supernatant was collected as the Unbound fraction, and IP sample beads were washed three times in IP Buffer. Finally, IP sample beads were resuspended in IP Buffer, transferred to clean tubes, and stored along with Input samples overnight at 4°C to allow passive hydrolysis to partially reverse formaldehyde crosslinks. This protocol was applied for both protein co-immunoprecipitation and RNA immunoprecipitation.

For protein co-immunoprecipitation, harsh elution of protein from IP sample beads was accomplished the next day—IP samples were diluted in modified Laemmli Sample Buffer (Laemmli 1970), incubated at 98°C for 5 minutes, centrifuged at 16,100×g for 5 minutes at room temperature, and magnetized to collect beads. Sample supernatants were then collected as IP samples. In parallel, Input samples were concentrated using an acetone-based method; this step was required for subsequent immunoblot analysis. Input samples were diluted to generate 80% chilled acetone solutions, vortexed for 15 seconds, and incubated at −20°C for 60 minutes. Samples were centrifuged at 14,000×g for 10 minutes at room temperature, resulting supernatants were discarded, and most remaining acetone was evaporated by air drying protein pellets in open tubes for 30 seconds at room temperature. To solubilize these dried protein pellets, samples were suspended in a solution equal parts modified Laemmli Sample Buffer (Laemmli 1970) and IP Buffer, vortexed, sonicated for 3×5 minutes in a 4°C Bioruptor ultrasonicator (UCD-200, Diagenode), vortexed, and heated at 98°C for 10 minutes. Finally, remaining insoluble material was collected by centrifugation at 16,100×g for 5 minutes at room temperature. Associated supernatants were isolated as concentrated Input protein samples. For RNA immunoprecipitation, harsh elution of RNA from IP sample beads was accomplished the next day with Trizol—both IP and Input samples were subjected to the RNA extraction protocol detailed below.

### RNA Extraction

Following immunoprecipitation, RNA was isolated from IP and Input samples using a TRIzol-column hybrid approach adapted from (Rodriguez-Lanetty). To account for volume differences, samples were vigorously homogenized in TRIzol reagent (15596018, Thermo Fisher) at a ratio of either 1:10 (IP sample:TRIzol) or 1:3 (Input sample:TRIzol) and then incubated for 5 minutes at room temperature. All homogenized samples were clarified by centrifugation at 12,000×g at 4°C for 5 minutes, IP samples were magnetized to collect beads, and supernatant was isolated from all samples. After adding chloroform at a ratio of 0.2:1 (choloroform:TRIzol), samples were manually shaken and incubated at room temperature for 3 minutes. Samples were phase separated by centrifugation at 12,000×g at 4°C for 15 minutes, after which the aqueous layer was carefully isolated and mixed with an equal volume of 100% ethanol. RNA was further purified using an RNeasy Mini Kit (74106, QIAGEN) according to the manufacturer’s instructions (RNeasy Mini Handbook, 4^th^ Ed., June 2012) with the following deviations: for each sample, a final 30 µL elution was performed twice, isolating 60 µL of RNA in total into each collection tube. An on-column DNase digestion step was also performed under the same instructions using an RNase-Free DNase Set (79254, QIAGEN). Final RNA concentration and sample purity were determined via a NanoDrop 1000 spectrophotometer (Thermo Fisher).

### RNA Sequencing

RNA from twelve samples of 300 adult female *Drosophila* heads each was isolated via the immunoprecipitation and extraction protocols described above, generating twelve pairs of IP and Input samples, or twenty-four samples in total. These samples were composed of four biological replicates each of *elav>Gal4* alone, *elav>Nab2-FLAG*, and *elav>Atx2-3xFLAG*. Once obtained, RNA samples were transferred on dry ice to the Georgia Genomics and Bioinformatics Core at UGA for library preparation and sequencing. There, IP samples were first concentrated using solid phase reversible immobilization (SPRI) beads. Then, the TruSeq Stranded Total RNA Library Prep Gold kit (20020598, Illumina) was used to deplete rRNA and prepare stranded cDNA libraries from all twenty-four samples. These uniquely barcoded cDNA libraries were then pooled by sample type, forming one IP library pool and one Input library pool. Each pool was sequenced on a separate NextSeq High Output Flow Cell (Illumina) for 150 cycles to generate paired-end, 75 base-pair (bp) reads. Total non-index sequencing yield across all IP samples was 88.49 Gbp, equivalent to about 1.2 billion reads in total and 98 million reads per sample. Total non-index sequencing yield across all Input samples was 83.25 Gbp, equivalent to about 1.1 billion reads in total and 93 million reads per sample. Sequencing accuracy was high; 87.83% and 91.38% of non-index reads for IP and Input samples, respectively, have a sequencing quality (Q) score greater than or equal to 30.

### RNA Sequencing Analysis—Read Mapping, Differential Expression, Visualization

Following sequencing, raw read FASTA files were transferred to Emory for bioinformatic analysis. To start, analyses were conducted on the Galaxy web platform, specifically using the public server at usegalaxy.org (Afgan *et al*. 2018). This analysis was supported by the BDGP6.22 release of the *Drosophila melanogaster* genome (Hoskins *et al*. 2015)—both the raw sequence FASTA and the gene annotation GTF were downloaded from release 97 of the Ensembl database (Yates *et al*. 2020) and used as inputs in subsequent read mapping, annotation, and visualization steps. For each Galaxy tool described below, exact parameters and version numbers used are detailed in Supplemental Table 1. For each sample, reads from across all four NextSeq flow cell lanes were concatenated using the Galaxy *Concatenate datasets tail-to-head* tool and mapped using RNA STAR (Dobin *et al*. 2013). Mapped reads were then assigned to exons/genes and tallied using *featureCounts* (Liao *et al*. 2014). To enable inter-sample read count comparisons, count normalization and differential expression analysis was conducted using *DESeq2* (Love *et al*. 2014). Importantly, *DESeq2* analysis was performed twice, once on the 12 IP samples and once on the 12 Input samples; see *Supplemental Materials and Methods* for discussion of this sample separation method.

Outputs from all of the above tools were downloaded from Galaxy for local analysis, computation, and visualization. Custom R scripts were written to generate the scatterplots and hypergeometric test reported here and are available in File S3. Scripts in the R programming language (R Core Team 2019) were written and compiled in RStudio (R Studio Team 2018). Additional R packages used in these scripts include ggplot2 (Wickham 2016), ggrepel (Slowikowski 2019), BiocManager (Morgan 2018), and DESeq2 (Love *et al*. 2014). Analyses were supported by bulk data downloads along with extensive gene-level annotation, sequence information, and references provided by Flybase (Thurmond *et al*. 2018). Principal component analysis was conducted by and reported from the above *DESeq2* assessment on Galaxy. Mapped reads were visualized in the Integrative Genomics Viewer (IGV) (Robinson *et al*. 2011) on the same version of the *D. melanogaster* genome used above.

### Gene-by-gene one-way ANOVAs to identify significantly enriched (i.e. RBP-associated) transcripts

Gene-by-gene ANOVAs and post-hoc tests for the 5,760 genes identified in the “testable” set, along with bar graphs of IP/Input values, were generated in Prism 8 for Windows 64-bit (GraphPad Software). Custom R and PRISM scripts were written to generate and label the 5,760 PRISM data tables, one per testable gene, required for this analysis, and custom R scripts were written to extract and combine the outputs from each test; these scripts are all available in File S3. See *Results* for a summary and below for a further detailed discussion of the statistical testing used to define the testable transcript set and identify significantly enriched (i.e. RBP-associated) transcripts in our RIP-Seq results.

To identify RNA targets of Nab2 and Atx2—that is, RNAs enriched in either Nab2 RIP or Atx2 RIP samples relative to control RIP—directly comparing normalized read counts between RIP samples is insufficient. Differences in RNA expression between samples must be accounted for, as these differences can partially or wholly explain differences in the amount of RNA isolated by IP. We employed a common solution to this problem used in RIP- and ChIP-qPCR (Zhao *et al*. 2010; Aguilo *et al*. 2015; Li *et al*. 2019), scaling normalized RIP reads for each gene in each sample by the corresponding number of normalized Input reads. For clarity, we describe these values as “IP/Input”— they are commonly referred to as “Percent Input” or “% Input.” These IP/Input values could then be compared between samples, further normalizing them to *elav-Gal4* alone controls. In this way, RIP fold enrichment, appropriately normalized to library size/composition *and* gene expression, were calculated for each gene in each sample. To promote the reliability of our analyses and increase our statistical power to detect differences in fold enrichment, we limited further analyses to a testable set of 5,760 genes out of the 17,753 total genes annotated in the BDGP6.22 genome. The testable gene set was defined as having detectable expression in all twelve Input samples and an average normalized read count in either Nab2 or Atx2 RIP samples greater than 10. These criteria were based on those used in (Lu *et al*. 2014; Malmevik *et al*. 2015). In this defined gene set, differences in fold enrichment were statistically tested using gene-by-gene one-way ANOVAs (Li *et al*. 2019) in Prism 8 (GraphPad software), applying Dunnett’s post-hoc test to calculate significance *p*-values only for the comparison of each experimental sample to the control sample (Dunnett 1955). In each case, *p*-values were adjusted to correct for multiple hypothesis testing only within each gene-by-gene ANOVA. We identified a small, focused set of statistically significantly enriched RNAs using this approach and concluded that additional corrections across all genes to control type I error (i.e. false positives) are not necessary (Rothman 1990). In fact, in the analyses above we determined that rRNA depletion during our RIP-Seq library preparation was incomplete, resulting in comparatively low read depth. Thus, rather than failing to adequately control type I error, we strongly suspect the RBP-associated transcripts we identified through this approach represent an undercount, to be expanded in future studies by methods with higher sensitivity (e.g. CLIP-Seq).

### RNA Sequencing Analysis—Sequence Motif Analyses

Sequence motif analyses were conducted using the MEME Suite of software tools, accessed through the web interface at meme-suite.org (Bailey *et al*. 2009). For each MEME Suite tool described below, exact parameters and version numbers used are detailed in Supplemental Table 1. Within the MEME Suite, we used MEME itself (Bailey and Elkan 1994) to scan all Nab2-associated transcripts, regardless of their association with Atx2, to 1) identify sequence motifs shared across multiple transcripts and 2) evaluate the frequency and statistical significance of the discovered sequence motifs. Next, FIMO (Grant *et al*. 2011) was used to quantify the frequency among 1) Nab2-associated transcripts and 2) non-Nab2 associated transcripts of user-provided sequences, specifically i) a 41-bp A-rich motif identified in Nab2-associated transcripts by MEME, ii) A_12_, and iii) A_11_G. Non-Nab2-associated transcripts are defined as all 5,619 transcripts in the testable set found to not be statistically significantly associated with Nab2 by RIP-Seq. Sequence logos (i.e. visual representations of weighted sequence motifs) were generated by MEME and by WebLogo 3.7.4, available at weblogo.threeplusone.com (Crooks *et al*. 2004).

Importantly, for any Nab2-associated or non-Nab2 associated transcripts annotated with multiple splice variants, all variant sequences were included as inputs in our motif analyses. This inclusion reflects an inherent limitation of standard shotgun—that is, short-read—sequencing, as most reads cannot be unambiguously assigned to one splice variant of a given gene, only to given exon(s) encoded by that gene. We therefore chose this inclusion strategy to avoid introducing any bias associated with attempting to call single splice variants for RBP association, and for analytical simplicity. Full sequences of Nab2-associated and non-Nab2 associated transcripts were obtained using the FlyBase Sequence Downloader at flybase.org/download/sequence/batch/ (database release FB2020_04).

### Data Availability

The authors affirm that all data necessary for confirming the conclusions of the article are present within the article and associated figures, tables, supplemental materials, and database accessions. File S1 contains *Supplemental Materials and Methods*, including those focused on bulk *Drosophila* head isolation, immunoblotting, *DESeq2*-based count normalization, and Gene Ontology analyses. File S2 contains detailed legends for all supplemental tables. File S3 contains all custom code—both R and PRISM scripts—written to generate, analyze, or visualize data in this article and associated figures, tables, and supplemental materials. Sequencing data, including raw reads, processed counts, and statistical analyses for each individual RIP-Seq sample, are available at the Gene Expression Omnibus (GEO) under accession: GSE165677. *Drosophila* stocks are available upon request. Supplemental materials, including files, figures, and tables, are available at figshare: https://figshare.com/s/6f28676d7119624b3105.

## Results

### *Atx2* loss-of-function alleles suppress Nab2 overexpression phenotypes in the adult eye

Previous work has established a Gal4-driven Nab2 overexpression system in the *Drosophila* eye as an effective screening platform to identify potential regulatory partners and targets of Nab2 (Pak *et al*. 2011; Bienkowski *et al*. 2017; Lee *et al*. 2020). This approach uses the *Glass Multimer Reporter* (*GMR*) construct (Ellis *et al*. 1993; Hay *et al*. 1994) to drive expression of the *S. cerevisiae* Gal4 transcription factor in fated eye cells (Freeman 1996). In turn, Gal4 binds to *Upstream Activating Sequence* (*UAS*) sites within an EP-type P-element (Rørth 1996) inserted upstream of the endogenous *Nab2* gene (*EP3716*) and induces eye-specific overexpression of endogenous Nab2 protein (a genotype hereafter referred to as *GMR>Nab2*). *GMR>Nab2* produces a consistent array of eye morphological defects compared to the *GMR-Gal4* transgene control (Pak *et al*. 2011; Bienkowski *et al*. 2017; Lee *et al*. 2020) and (Figure 1A,B). Specifically, this misexpression causes loss of posterior eye pigment, sporadic blackened patches, and disruptions to ommatidial organization lending the surface of the eye a “rough” appearance. Notably, *GMR>Nab2*-induced pigment loss increases in severity along the anterior-to-posterior axis of the eye, likely because *GMR* activation occurs behind the morphogenetic furrow, the posterior-to-anterior wave of eye morphogenesis observed in the larval eye disc (Wolff and Ready 1991; Hay *et al*. 1994). As a result, posterior *GMR>Nab2* eye cells experience the longest period of Nab2 overexpression.

**Figure 1.**
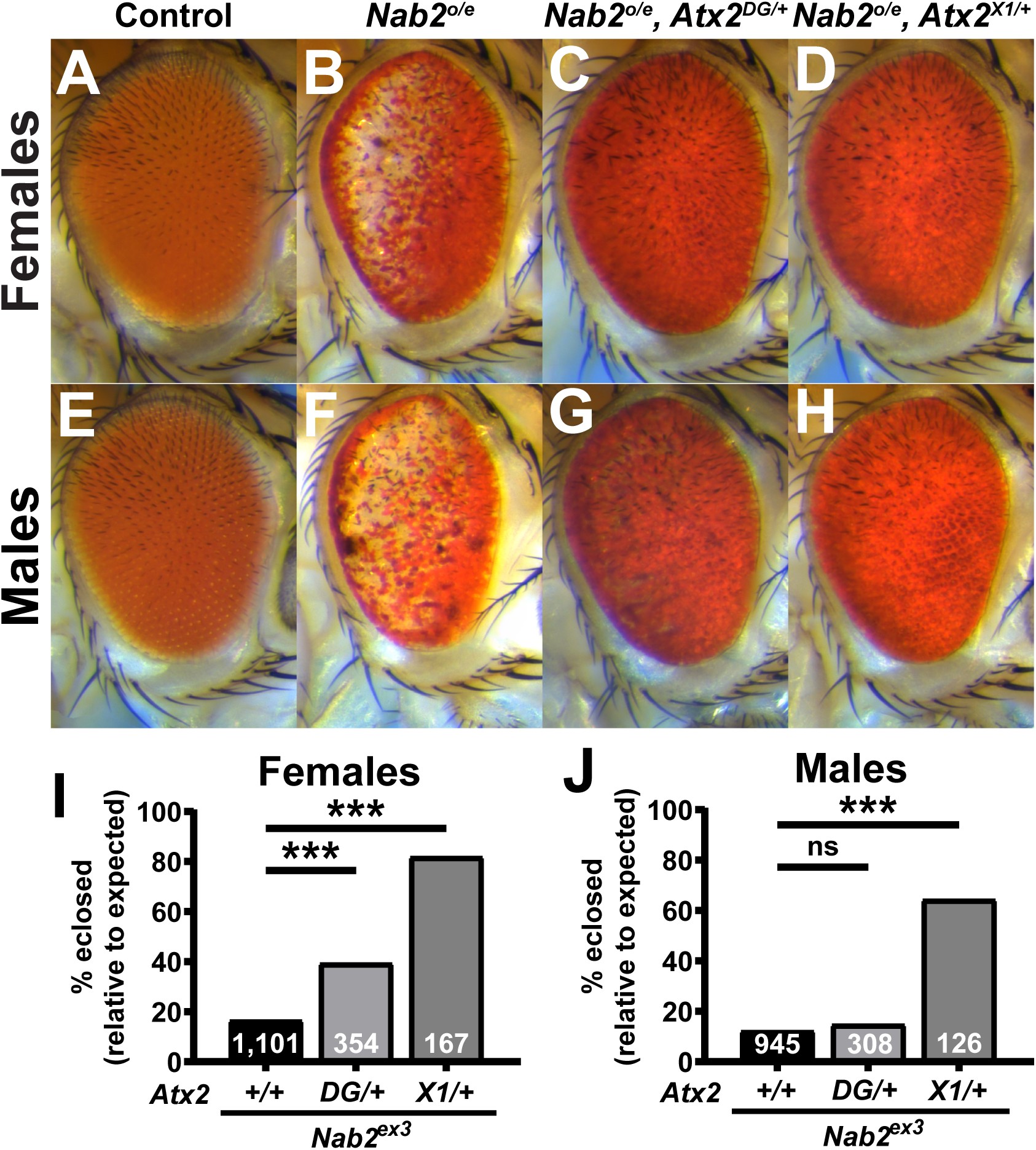
Loss-of-function alleles of *Atx2* suppress effects of *Nab2* misexpression in female and male *Drosophila*. Compared to (A) the uniform color and regimented ommatidial structure of the *Drosophila* eye in control females expressing the fated-eye-cell-specific *GMR-Gal4* driver alone, (B) overexpression of endogenous *Nab2* with *GMR-Gal4* (*Nab2^o/e^*) induces posterior pigment loss, sporadic blackened patches, and ommatidial disorder or “roughness”. Heterozygosity for either of two *Atx2* loss-of-function alleles, (C) *Atx2^DG08112/+^* or (D) *Atx2^X1/+^*, dominantly suppresses the pigment loss and blackened patch phenotype, with limited impact on roughness. (E-H) These genetic relationships are also observed in eyes in males. (I, J) Flies lacking functional endogenous *Nab2*, *Nab2^ex3^* homozygotes, demonstrate dramatically decreased adult viability, as quantified by the percentage of flies reaching pupal eclosion and adulthood out of the amount expected by Mendelian inheritance. (I) In females, both loss-of-function alleles of *Atx2* significantly suppress this effect, partially rescuing viability; (J) in males, only *Atx2^X1/+^* suppresses. Sample sizes (n) are reported in each bar and include all F1 progeny scored, including genetically distinct siblings of the genotype of interest used to calculate *% eclosed (relative to expected)*. Fisher’s Exact Test (two-sided) was used to assess statistical significance. ns=not significant, ***=*p*<0.001.

Using the *GMR>Nab2* modifier screen as a foundation, we previously identified the *Drosophila* Fragile X Syndrome RBP and neuronal translational regulator Fmr1 as a physical and functional interactor of Nab2 (Bienkowski *et al*. 2017). An allele of the *Ataxin-2* (*Atx2*) gene, which encodes an RNA binding protein that is a regulatory partner of Fmr1 in *Drosophila* (Sudhakaran *et al*. 2014), was also detected in eye this screen as a candidate *GMR>Nab2* modifier (Bienkowski *et al*. 2017). To pursue this potential Nab2-Atx2 link, we tested two *Atx2* alleles for genetic interactions with *GMR>Nab2*. The first allele, *Atx2^DG08112^*, is caused by the insertion of a 15.6 kb *{wHy}* P-element near the 5’ end of *Atx2* (Huet *et al*. 2002; Bellen *et al*. 2004) and is lethal *in trans* to *Df(3R)Exel6174*, a deletion that completely removes the *Atx2* locus and nearby genes (Parks *et al*. 2004). That is, crossing balanced *Atx2^DG08112^* and *Df(3R)Exel6174* alleles produces no *trans* heterozygotes among other F1 progeny (n=54). Based on these data, we interpret *Atx2^DG08112^* to be a strong hypomorph. The second *Atx2* allele, *Atx2^X1^*, is a 1.4 kb imprecise-excision-based deletion that removes the first 22 codons of the *Atx2* coding sequence and that has been characterized as a null (Satterfield *et al*. 2002). In part because Nab2 loss induces some sex-specific defects (Jalloh *et al*. 2020), we analyzed each sex individually. In adult females, heterozygosity for either of these two loss-of-function alleles, *Atx2^DG08112^* (Figure 1C) or *Atx2^X1^* (Figure 1D), dominantly suppresses the pigment loss and blackened patches caused by *GMR>Nab2*. In contrast, both *Atx2* alleles have limited impact on ommatidial organization or “roughness”. In males, *GMR>Nab2* induces morphological eye defects (Figure 1 E,F) comparable to those in females, and similarly heterozygosity for either *Atx2^DG08112^* (Figure 1G) or *Atx2^X1^* (Figure 1H) dominantly suppresses the pigment loss and blackened patch defects.

### *Atx2* loss-of-function alleles suppress *Nab2* null effects on adult viability and brain morphology

Misexpression of Nab2 induces dramatic phenotypes in domains beyond the eye; homozygosity for the null allele *Nab2^ex3^* causes a dramatic reduction in adult viability (Pak *et al*. 2011). Thus, to explore whether modifying effects of *Atx2* loss-of-function alleles extend to the endogenous *Nab2* locus, we analyzed the effect of *Atx2* heterozygosity on low adult viability in *Nab2^ex3^* homozygotes (Supplemental Figure 1). As in the eye, both the *Atx2^DG08112^* and *Atx2^X1^* alleles dominantly suppress the viability defects observed in *Nab2^ex3^* females, elevating adult viability from 17% to 39% and 82%, respectively (Figure 1I). The corresponding effect in males is not as penetrant; only the null *Atx2^X1^* allele dominantly suppresses the viability defect in *Nab2^ex3^* males (Figure 1J). Taken together, these data establish gross similarities in *Nab2*-*Atx2* genetic interactions in females and males. Thus, for practicality we focused further experiments exclusively on female flies, given the more prohibitive impact on male viability of changes in Nab2 expression (Jalloh *et al*. 2020 and see *Materials and Methods*).

That *Atx2* loss-of-function alleles improve adult viability of *Nab2^ex3^* homozygotes suggests Atx2 and Nab2 coregulate processes or transcripts important for adult development or survival. However, these genetic interactions do not reveal in what cell types or tissues this coregulation may occur. We therefore focused further investigations of *Nab2*-*Atx2* interaction in the brain, given the established and important roles of each protein in brain neurons (Lim and Allada 2013; Sudhakaran *et al*. 2014; Kelly *et al*. 2016; Bienkowski *et al*. 2017). *Nab2^ex3^* homozygous flies develop morphological defects in the axon tracts—lobes—of the mushroom body (MB) brain structure, a principal olfactory learning and memory center of the insect brain (Heisenberg 2003; Kahsai and Zars 2011; Yagi *et al*. 2016; Takemura *et al*. 2017). Specifically, the MBs of surviving *Nab2^ex3^* homozygous null adults exhibit two highly penetrant structural defects: thinning or absence of the dorsally-projecting α lobes and over-projection or “fusion” of the medially-projecting β lobes (Kelly *et al*. 2016). We found that heterozygosity for either *Atx2^DG08112^* or *Atx2^X1^* also causes defects in MB morphology—specifically β lobe fusion—with no apparent effects on α lobe morphology as compared to controls (Figure 2A-C). Importantly, in the background of *Nab2^ex3^* nulls (Figure 2D), heterozygosity for either *Atx2^DG08112^* (Figure 2E) or *Atx2^X1^* (Figure 2F) suppresses the thinning or absence of α lobes, decreasing the penetrance of this phenotype from 62% of α lobes to 30% or 36%, respectively (Figure 2G). In contrast, neither *Atx2* allele significantly affects the penetrance of β lobe fusion in *Nab2^ex3^* nulls, demonstrating the effect of each mutation is not additive to the effect of *Nab2^ex3^* homozygosity in this context (Figure 2H). A similar α-lobe-specific interaction occurs between alleles of *Nab2* and *Fmr1* (Bienkowski *et al*. 2017). Notably, as α and β lobes are composed of tracts of bifurcated axons from single cells (Takemura *et al*. 2017), this α-lobe-specific suppression by *Atx2* alleles demonstrates a *Nab2*-*Atx2* genetic interaction at subcellular resolution. Moreover, that *Atx2* loss-of-function alleles suppress defects of a *Nab2* null allele implies that Atx2 and Nab2 proteins may coregulate, but in opposing ways, pathways guiding α lobe morphology during development.

**Figure 2.**
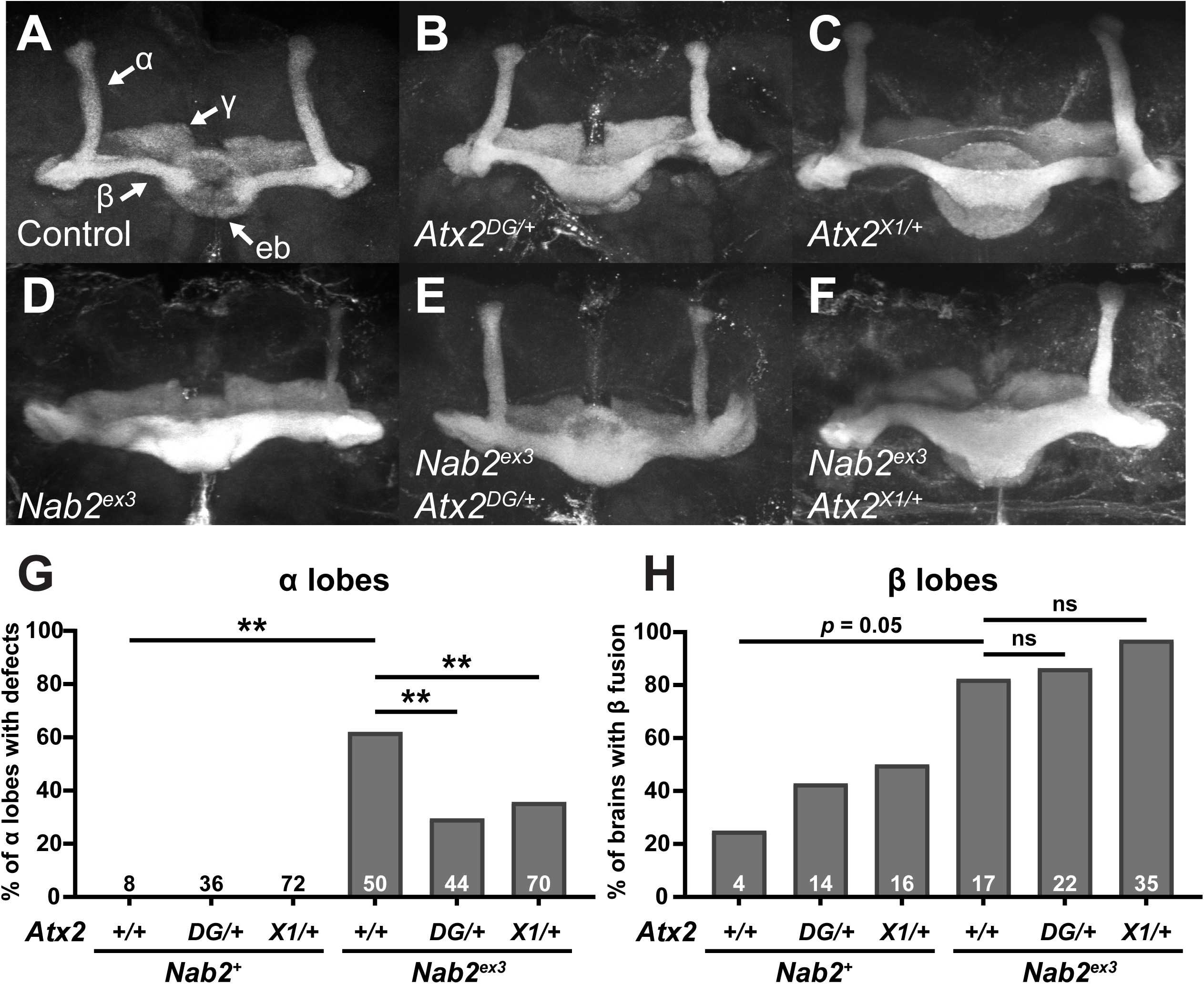
Loss-of-function alleles of *Atx2* specifically suppress axon morphology defects in *Nab2^ex3^* mushroom body α, but not β, lobes. (A) In a representative *Nab2^pex41^* control brain, Fasciclin 2 (Fas2)-marked axons from some Kenyon cells of the mushroom body bifurcate and project dorsally into α lobes and medially into β lobes. Fas2 also marks mushroom body γ lobes and the ellipsoid body (eb) (white arrows). Representative images show heterozygosity for (B) *Atx2^DG08112/+^* or (C) *Atx2^X1/+^* induces over-projection or “fusion” of β lobes, while (D) homozygosity for the *Nab2* null allele *Nab2^ex3^* induces both β lobe fusion and the thinning or complete absence of α lobes. Heterozygosity for either (E) *Atx2^DG08112/+^* or (F) *Atx2^X1/+^* in combination with *Nab2^ex3^* partially restores proper α lobe morphology and, as quantified in (G), significantly suppresses the penetrance of α lobe defects compared to *Nab2^ex3^* alone. (H) By comparison, as quantified in (H), these *Atx2* alleles neither suppress nor enhance the penetrance of β lobe defects compared to *Nab2^ex3^* alone. Sample sizes (n) are reported in each bar and quantify, for each genotype, the total number of α lobes scored for defects and the total number of brains scored for β lobe fusion. Fisher’s Exact Test (two-tailed) was used to assess statistical significance. ns=not significant, **=*p*≤0.01.

### Nab2 and Atx2 primarily localize to independent compartments in mushroom body neurons

The genetic links between *Nab2* and *Atx2* could reflect a physical interaction between their encoded proteins (e.g. as shared components of mRNP complexes), as has been observed for both Nab2 and Atx2 with Fmr1 (Sudhakaran *et al*. 2014; Bienkowski *et al*. 2017). Alternatively, these genetic links could reflect functional but not physical interactions between Nab2 and Atx2 on common RNAs or neurodevelopmental processes. The latter hypothesis aligns with the localization patterns of each protein—Nab2 localizes primarily to neuronal nuclei with a small fraction in the cytoplasm in some contexts (Kelly *et al*. 2016; Bienkowski *et al*. 2017), while Atx2 is exclusive to the neuronal cytoplasm except under certain pathogenic conditions (Lessing and Bonini 2008; Elden *et al*. 2010). To begin to differentiate between these hypotheses, we evaluated the localization profiles of each protein in MBs *in vivo*. We expressed both *UAS-Nab2-YFP* and *UAS-Atx2-3xFLAG* transgenes in adult MB Kenyon cells using the pan-MB driver *OK107-Gal4* (Figure 3A). Similar to observations in human cerebral cortex tissues (Huynh *et al*. 2003), Atx2 is nearly excluded from nuclei and localizes strongly to the soma cytoplasm of MB Kenyon cells in adults *in vivo*. In contrast, Nab2 localizes predominantly to the nuclei of these neurons *in vivo*. This distinction extends beyond the soma and into the α and β lobe axon tracts; Atx2 localizes robustly to these cytoplasmic compartments while Nab2 does not (Supplemental Figure 2).

**Figure 3.**
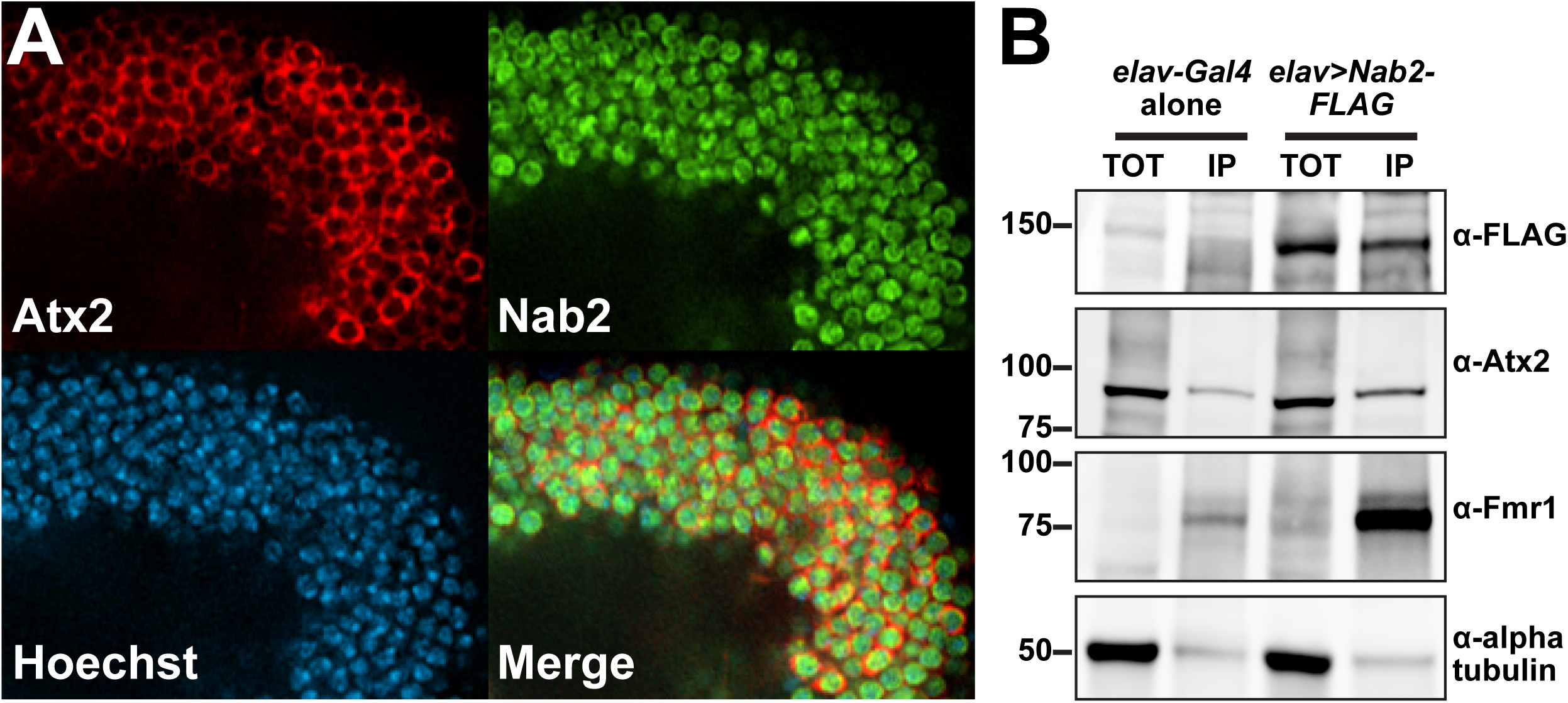
Nab2 and Atx2 primarily localize to different cellular compartments and show limited physical association in brain neurons. (A) To specifically assess protein localization in mushroom body neurons, tagged transgenic copies of Atx2 and Nab2 (Atx2-3xFLAG and Nab2-YFP) were expressed in female brains under the MB-specific *OK107-Gal4*. Kenyon cell soma, the cell bodies of the MBs, are shown for a representative brain. False-colored panels show fluorescence corresponding to α-FLAG (red, Atx2-3xFLAG), α-GFP (green, Nab2-YFP), Hoechst 33342 (blue, nuclei), and a merge of all three channels. Nab2 is localized primarily to the nuclei at steady state based on overlap with Hoechst 33342 signal, and Atx2 localizes primarily in the surrounding cytoplasm. (B) To test for physical association between Nab2 and Atx2 in brain neurons, lysates of female *Drosophila* heads, either *elav-Gal4* alone controls or *elav>Nab2-FLAG*, were subjected to co-immunoprecipitation using α-FLAG. For both genotypes, Input samples (TOT) represent 6.25% of assayed lysate and immunoprecipitation (IP) samples represent 25% of total samples eluted from α-FLAG beads. Samples were resolved via gel electrophoresis and analyzed by immunoblotting, probing with antibodies against FLAG, Atx2, Fmr1 (a positive control), or alpha tubulin (a negative control). Atx2 associates weakly with Nab2 based on its enrichment in IP samples; this association is less robust than that between Nab2 and positive control Fmr1.

To more rigorously assess Nab2-Atx2 protein interactions across all cell compartments, we expressed a FLAG-tagged Nab2 transgene (*UAS-Nab2-FLAG*) (Pak *et al*. 2011) using the pan-neuronal driver *elav-Gal4* (Lin and Goodman 1994) and subjected brain-neuron-enriched head lysates to immunoprecipitation with α-FLAG-conjugated beads to recover Nab2-associated proteins. Probing with specific antibodies confirms that Fmr1 is enriched in Nab2 immunoprecipitates as previously reported (Bienkowski *et al*. 2017), but reveals weak enrichment of Atx2 (Figure 3B). These results indicate complexes containing Nab2 and Atx2 may form in neurons but are rare relative to Nab2-Fmr1 complexes. Taken together, these subcellular localization and biochemical data suggest Nab2 and Atx2 do not generally co-occupy the same RNA or mRNP complexes throughout the post-transcriptional life of an RNA in adult mushroom body neurons. Therefore, we considered the possibility that *Nab2*-*Atx2* genetic interactions instead reflect roles in post-transcriptional control of shared RNA targets at different points in time or different locations in the cell.

### The Nab2 and Atx2 RNA interactomes in brain neurons overlap

Neither Nab2-nor Atx2-associated RNAs have been identified by a high-throughput method in *Drosophila*—such accounting has been conducted for Atx2 in human cells (Yokoshi *et al*. 2014) and for Nab2 only in *S. cerevisiae*, not in any metazoan (Guisbert *et al*. 2005; Batisse *et al*. 2009; Tuck and Tollervey 2013; Baejen *et al*. 2014). To test the hypothesis that Nab2 and Atx2 share RNA targets, we identified transcripts stably associated with epitope-tagged versions of each protein in adult brain neurons using an RNA immunoprecipitation-sequencing (RIP-Seq) approach. In this approach, protein products of *UAS-Nab2-FLAG* or *UAS-Atx2-3xFLAG* transgenes are robustly expressed under *elav-Gal4* control and are efficiently immunoprecipitated from adult head lysates (Figure 4A). Briefly, four biological replicates each of *elav-Gal4*, *elav>Nab2-FLAG*, and *elav>Atx2-3xFLAG* adult female *Drosophila* heads were lysed and immunoprecipitated with α-FLAG-conjugated beads. Then, RNA from both IP and Input samples was rRNA depleted, reverse transcribed into stranded cDNA libraries, and sequenced. Using the Galaxy web platform through the public server at usegalaxy.org (Afgan *et al*. 2018), reads were mapped using STAR (Dobin *et al*. 2013) to the BDGP6.22 release of the *Drosophila melanogaster* genome (sourced through Ensembl, Yates *et al*. 2020), assigned to exons/genes and tallied using *featureCounts* (Liao *et al*. 2014), and normalized for inter-library count comparisons using *DESeq2* (Love *et al*. 2014). A principal component analysis (PCA) generated as part of *DESeq2* demonstrates the high inter-genotype reproducibility among RNA IP (RIP) samples and shows that samples expressing Nab2-FLAG or Atx2-3xFLAG differ more from *elav-Gal4* controls than from one another (Figure 4B).

**Figure 4.**
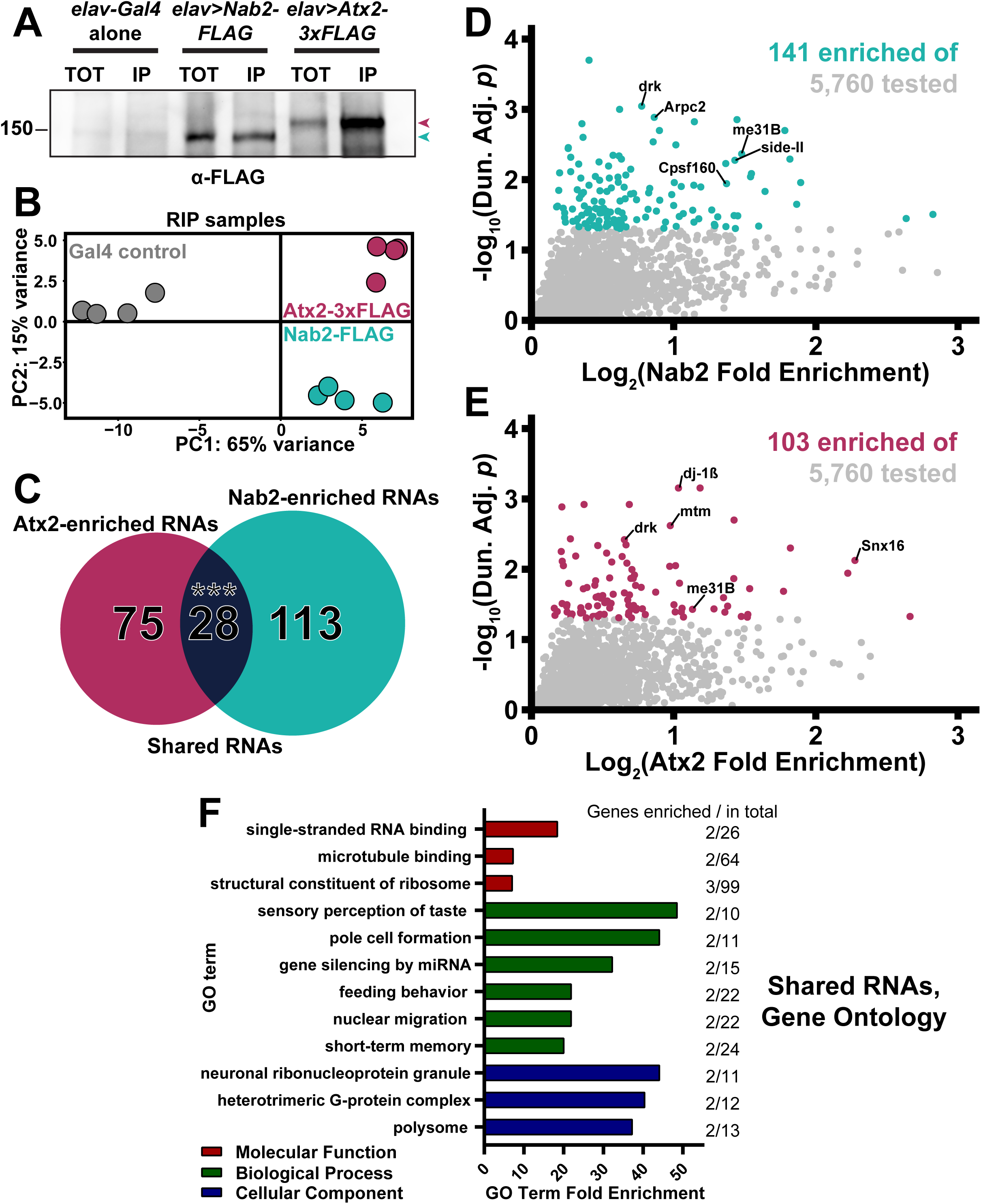
RIP-Seq reveals overlapping sets of transcripts associate with Nab2-FLAG and Atx2-3xFLAG in brain neurons. (A) Lysates from heads of female adult flies expressing either pan-neuronal *elav-Gal4* alone as a control, *elav>Nab2-FLAG*, or *elav>Atx2-3xFLAG* were subjected to α-FLAG immunoprecipitation and immunoblotting to test IP efficacy. Input samples (TOT) represent ∼6.25% of total assayed lysates and immunoprecipitation samples (IP) represent 25% of total samples eluted from α-FLAG beads. Both epitope-tag samples show robust immunoreactivity to α-FLAG in TOT and IP (arrowheads), indicating effective transgene expression and successful tagged-protein enrichment by IP. (B) Principal component analysis of 12 sequenced RNA IP samples reveals high intra-genotype reproducibility. Comparison of principal component 1 (PC1) and principal component 2 (PC2) demonstrates Nab-FLAG (teal) and Atx2-3xFLAG (maroon) samples differ more from Gal4 controls (gray) than from one another, as predicted. (C) Venn diagram of Nab2-enriched and Atx2-enriched RNAs identified by RIP-Seq, revealing that 28 shared transcripts associate with both RBPs, a significant overlap according to the hypergeometric test (***=*p*<0.001). (D-E) Scatter plot of all transcripts within the 5,760 of the testable set with positive (D) *log_2_(Nab2 Fold Enrichment)* or (E) *log_2_(Atx2 Fold Enrichment)* values. *Fold Enrichment* values quantify how effectively a transcript was enriched by IP and are derived by calculating IP/Input (i.e. percent input) values for control and epitope-tag samples and setting the average of control values to 1 (i.e. 0 on the logarithmic scale used here). Y-axes display results of significance testing, conducted by gene-by-gene one-way ANOVA, Dunnett’s post-hoc test, and within-gene multiple hypothesis testing adjustment (*Dun. Adj. p*). Statistically significant transcripts (*Dun. Adj. p* < 0.05) are colored. On each plot, labels identify three transcripts among the “top” (see *Results* for details) RBP-specific RBP-associated transcripts and two transcripts (*drk*, *me31B*) among the shared RBP-associated transcripts. (F) The independent *Molecular Function* (red), *Biological Process* (green), and *Cellular Component* (blue) Gene Ontology (GO) terms most overrepresented among the shared Nab2-and Atx2-associated transcripts as compared to the entire testable transcript set. GO term independence was determined by “Hierarchical Selection” (see *Methods*). The number of GO term members within the shared RBP-associated transcripts and within the entire testable transcript set (*Genes enriched / in total*) are reported to the right of each bar.

To identify Nab2-associated and Atx2-associated RNAs, we calculated percent input (IP/Input) enrichment values (Zhao *et al*. 2010; Aguilo *et al*. 2015; Li *et al*. 2019) for each of the 5,760 genes in the testable set defined by 1) detectable expression in all twelve Inputs and 2) an average normalized Nab2-or Atx2-IP read count greater than 10 (Lu *et al*. 2014; Malmevik *et al*. 2015). Fold enrichment differences were statistically tested by performing gene-by-gene one-way ANOVAs (Li *et al*. 2019), applying Dunnett’s post-hoc test (Dunnett 1955), and calculating adjusted *p*-values corrected for multiple hypothesis testing within each gene-by-gene ANOVA (values hereafter referred to as *Dun. Adj. p*; see *Materials and Methods* for more detail). Using this approach, we identify 141 and 103 RNAs significantly enriched in α-FLAG IPs of *elav>Nab2-FLAG* and *elav>Atx2-3xFLAG* female heads, respectively (Supplemental Table 2, Supplemental Figure 3). The size and focus of these sets of statistically significantly enriched RNAs suggests type I (i.e. false positive) error is sufficiently controlled and additional corrections between genes are not necessary (Rothman 1990). Comparing the Nab2- and Atx2-IP groups strongly supports our hypothesis, revealing 28 transcripts shared between Nab2- and Atx2-associated *Drosophila* neuronal RNAs (Figure 4C). This overlap is highly significant according to the hypergeometric test—it is extremely unlikely to occur by random selection from the total tested gene set. The full list of transcripts associated with both Nab2 and Atx2 (Table 1) includes multiple mRNAs that encode proteins with functions in neuronal domains in which *Nab2* and *Atx2* genetically interact, raising the possibility that coregulation of these RNAs by Nab2 and Atx2 partially explains these *Nab2*-*Atx2* genetic links. These shared transcripts include *drk* (*downstream of receptor kinase*), *me31B* (*maternal expression at 31B*), *sm* (*smooth*), and *stai* (*stathmin*). The protein encoded by *drk* is a receptor tyrosine kinase (RTK) adaptor that regulates growth and development by binding activated RTKs, such as sevenless in R7 retinal cells (Almudi *et al*. 2010), and contributes to, among other processes, cell survival in the eye (Schoenherr *et al*. 2012) and olfactory learning and memory in the MB (Moressis *et al*. 2009). The protein encoded by *me31B* is a DEAD-box RNA helicase expressed in many cellular contexts, including the MB Kenyon cells (Hillebrand *et al*. 2010) and the oocyte (Nakamura *et al*. 2001), that physically associates with Atx2 (Lee *et al*. 2017) and serves as a central player in miRNA-mediated translational repression (Barbee *et al*. 2006) and assembly of some RNP granules (Eulalio *et al*. 2007). Finally, the proteins encoded by *sm* and *stai* are respectively an hnRNP linked to defects in axon termination (Layalle *et al*. 2005) and a tubulin binding protein linked to natural variation in the size of MB α and β lobes (Lachkar *et al*. 2010; Zwarts *et al*. 2015).

**TABLE 1.**
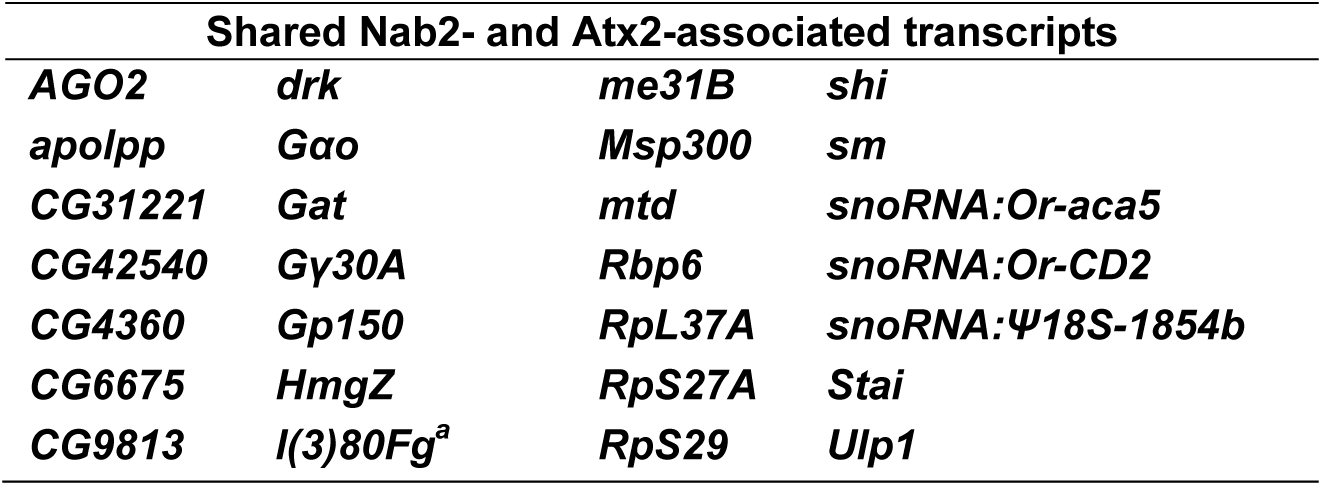
Identities of the 28 transcripts overlapping between the Nab2 and Atx2 RNA interactomes. For all 5,760 genes in the RIP-Seq testable set, control-normalized IP/Input enrichment values were calculated followed by gene-by-gene one-way ANOVAs, Dunnett’s post-hoc tests, and within-gene multiple hypothesis testing adjustment (*Dun. Adj. p*). All transcripts statistically significantly (*Dun. Adj. p* < 0.05) enriched in both Nab2- and Atx2-associated transcripts sets are listed here. Functional interactions between Nab2 and Atx2 in brain neurons may be explained by their coordinate regulation of these shared associated transcripts. *^a^*Symbol updated from *CG40178* to current nomenclature in BDGP6.37.

The 28 shared transcripts represent approximately 20% and 24% of the total transcripts identified as Nab2- and Atx2-associated, respectively, underscoring that these proteins also associate with RNA sets independent from one another. From these independent sets, we defined the top Nab2-specific and Atx2-specific associated transcripts as the top 20 most significantly associated transcripts (by *Dun. Adj. p*) and top 20 most strongly enriched transcripts (by IP/Input) in each set. As with shared RNAs, multiple RBP-specific RNAs with links to *Nab2* or *Atx2* functions or mutant phenotypes are identified among these top transcripts, raising the possibility that regulation of these RNAs by Nab2 or Atx2 partially explains the mechanism of action of these RBPs (Figure 4D,E). For example, the top Nab2-specific associated RNAs include *Arpc2* (*Actin-related protein 2/3 complex, subunit 2*), *side-II* (*sidestep II*), and *Cpsf160* (C*leavage and polyadenylation specificity factor 160*). These transcripts respectively encode proteins with proposed functions in neuronal growth cone advance (Yang *et al*. 2012), synapse formation between certain neuronal subtypes (Tan *et al*. 2015), and mRNA poly(A)-tail formation based on orthology to mammalian *Cpsf1* (Mandel *et al*. 2008). The top Atx2-specific associated RNAs include *dj-1β*, *mtm* (*myotubularin*), and *Snx16* (*Sorting nexin 16*). These transcripts respectively encode proteins with proposed functions in ATP synthesis and motor neuron synaptic transmission (Hao *et al*. 2010; Oswald *et al*. 2018), endosomal trafficking regulation via phosphatase activity (Velichkova *et al*. 2010; Jean *et al*. 2012), and neuromuscular junction synaptic growth (Rodal *et al*. 2011).

### Gene Ontology terms enriched in Nab2 and Atx2 RNA interactomes emphasize additional RBP-associated transcripts

Evaluating Nab2- and Atx2-associated RNAs individually provides valuable but incomplete insight, allowing larger trends to be missed. To complement these analyses, we holistically evaluated the shared and specific Nab2- and Atx2-associated transcripts by subjecting each gene list to PANTHER Gene Ontology (GO) analysis, revealing the identities and members of enriched GO terms in each transcript set (Ashburner *et al*. 2000; Mi *et al*. 2019; The Gene Ontology Consortium 2019). Critically, GO term enrichment was calculated by comparing term abundance between these lists and the testable set of 5,760 head-enriched genes rather than the entire genome. In this way, these analyses did not identify GO terms as enriched simply because of their overrepresentation in *Drosophila* heads. Among shared Nab2- and Atx2-associated RNAs, we identify overrepresented GO terms and RBP-associated transcripts within them that highlight crucial functions and processes Nab2 and Atx2 may coregulate (Figure 4F). Among these GO terms are ‘microtubule binding’, which includes *apolpp* (*apolipophorin*) and *shi* (*shibire*); ‘sensory perception of taste’, which includes *Gαo* and *Gγ30A*; ‘gene silencing by miRNA’, which includes *AGO2* (*Argonaute 2*) and *me31B*; and ‘short-term memory’, which includes *shi* and *drk*. Survey of the associated RNAs specific to either RBP reveals overrepresented GO terms and transcripts within them which may mediate processes Nab2 and Atx2 regulate independently of one another, including respectively the GO terms ‘exosomal secretion’, which includes *Rab35* and *Rab7*; and ‘regulation of ATP metabolic process’, which includes *Dg* (*Dystroglycan*) and *dj-1β* (Supplemental Figure 4).

To combine and summarize the individual transcript and GO analyses, we highlight groups of seven transcripts found within the shared (Figure 5A) and RBP-specific (Figure 5B,C) associated transcript sets. These highlights were selected from the combined set of transcripts 1) demonstrating a fold enrichment (IP/Input) greater than 1.5 and/or 2) included in the most overrepresented GO terms (fully defined in Supplemental Table 3). Beyond transcripts already described, this summary includes the shared transcript *HmgZ* (*HMG protein Z*), Nab2-specific transcripts *fwe* (*flower*) and *SLC22A* (*SLC22A family member*), and Atx2-specific transcripts *tea* (*telomere ends associated*) and *Xpc* (*Xeroderma pigmentosum, complementation group C*). A group of functionally diverse transcripts in the testable set that did not associate with either RBP is shown for comparison and as evidence of the specificity of the RIP-Seq assay (Figure 5D).

**Figure 5.**
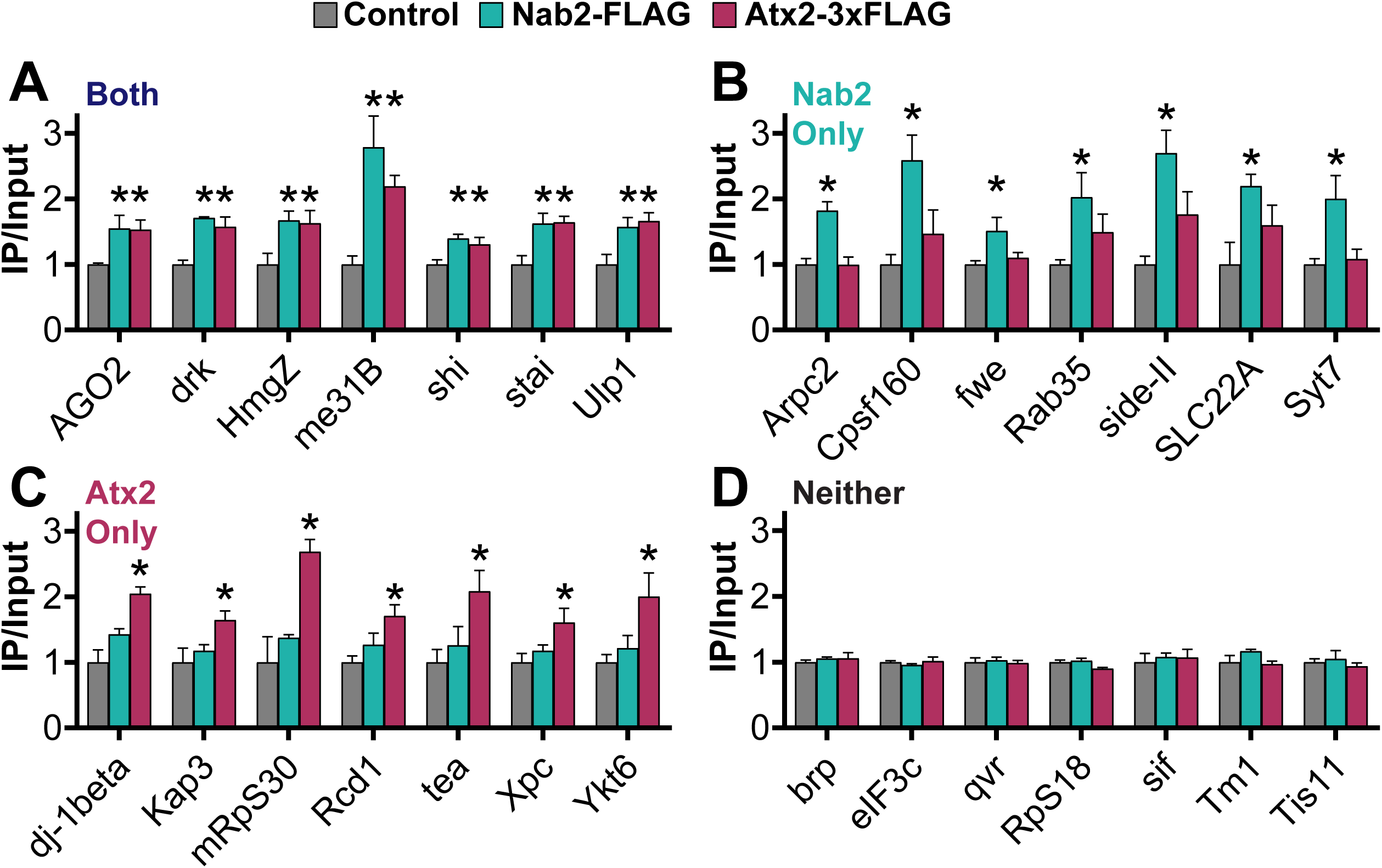
Potential functionally important RNA targets of Nab2 and Atx2 identified by combining individual transcript and holistic GO analyses of RIP-Seq results. For transcripts that associate with *Both* Nab2 and Atx2, *Nab2 Only*, *Atx2 Only*, or *Neither* RBP by RIP-Seq, seven transcripts of particular functional interest are presented as a summary of each category. (A-C) These transcripts met one or both of two criteria: 1) inclusion in an associated overrepresented GO term 2) an *IP/Input* (i.e. *Fold Enrichment*) value > 1.5. Given the functions of proteins encoded by these transcripts, these selections represent potential phenotypically important targets of post-transcriptional regulation by Nab2 and Atx2. (D) These transcripts, as a negative control, encode a functionally diverse set of proteins and do not associate with Nab2 or Atx2 (*Neither*), affirming the specificity of the RNA interactome of each RBP. Error bars represent standard errors of the mean (SEM). Gene-by-gene one-way ANOVA, Dunnett’s post-hoc test, and within-gene multiple hypothesis testing adjustment (*Dun. Adj. p*) was used to assess statistical significance. * = *Dun. Adj. p* < 0.05.

### Polyadenosine sequence motifs are enriched in Nab2-associated RNAs

The diversity of RNAs that do not associate with Nab2 and Atx2 in the RIP assay (Figure 5D) underscores a key finding—both of these RBPs exhibit specific RNA-association patterns within brain neurons. This observation is not surprising for Atx2 given, for example, the sequence specificity of its human homolog in HEK293T cells (Yokoshi *et al*. 2014), but it represents a valuable insight for Nab2. The extent of the metazoan Nab2/ZC3H14 RNA target pool has been an enduring question (Rha *et al*. 2017a), given the breadth of the *S. cerevisiae* Nab2 target pool (Batisse *et al*. 2009; Tuck and Tollervey 2013) and the ability of Nab2/ZC3H14 across eukaryotes to bind polyadenosine RNA *in vitro* (Kelly *et al*. 2007; Pak *et al*. 2011), raising the possibility for very broad binding of mRNAs via their poly(A) tails *in vivo*. We found a relatively focused set of RNAs co-precipitate with Nab2-FLAG from fly brain neurons, indicating Nab2 may indeed exhibit greater specificity in *Drosophila* than would be observed if the protein bound all or most polyadenylated transcripts via their poly(A) tails.

Thus, we sought to determine what additional RNA sequence features may drive the association of Nab2 with its target transcripts if not only the presence of a poly(A) tail. We used MEME (Bailey and Elkan 1994) to scan all Nab2-associated transcripts to identify shared sequence motifs that may represent Nab2 binding sites and partially explain Nab2 specificity. Strikingly, this analysis identifies a 41-bp long, internal-A-rich stretch among the first ten 6-50-bp motifs shared among Nab2-associated transcripts. Importantly, each of these 10 sequence motifs are shared across overlapping sets of many but not all Nab2-associated RNAs. Using FIMO (Grant *et al*. 2011), another part of the MEME Suite (Bailey *et al*. 2009), we quantified the frequency of close and exact matches to the consensus version of this motif among Nab2-associated RNAs. Occurrences of this A-rich motif are significantly more common in Nab2-associated transcripts compared to non-Nab2 associated transcripts, respectively appearing once every 135 bases and once every 845 bases on average, a 6.3-fold enrichment (Figure 6A). The high frequency of this motif in Nab2-associated transcripts is consistent with data from *S. cerevisiae* that Nab2 does not associate with RNAs exclusively through the poly(A) tail and also binds to upstream UTRs and coding sequences, likely through other A-rich sequences (Guisbert *et al*. 2005; González-Aguilera *et al*. 2011; Tuck and Tollervey 2013; Baejen *et al*. 2014; Aibara *et al*. 2017). Importantly, that this A-rich motif is enriched in but not exclusive to Nab2-associated RNAs is consistent with results for other RBPs—linear sequence motifs alone are generally insufficient to explain RBP specificity (Dominguez *et al*. 2018) and RBPs do not generally occupy all of their available binding motifs throughout the transcriptome (Li *et al*. 2010; Taliaferro *et al*. 2016).

**Figure 6.**
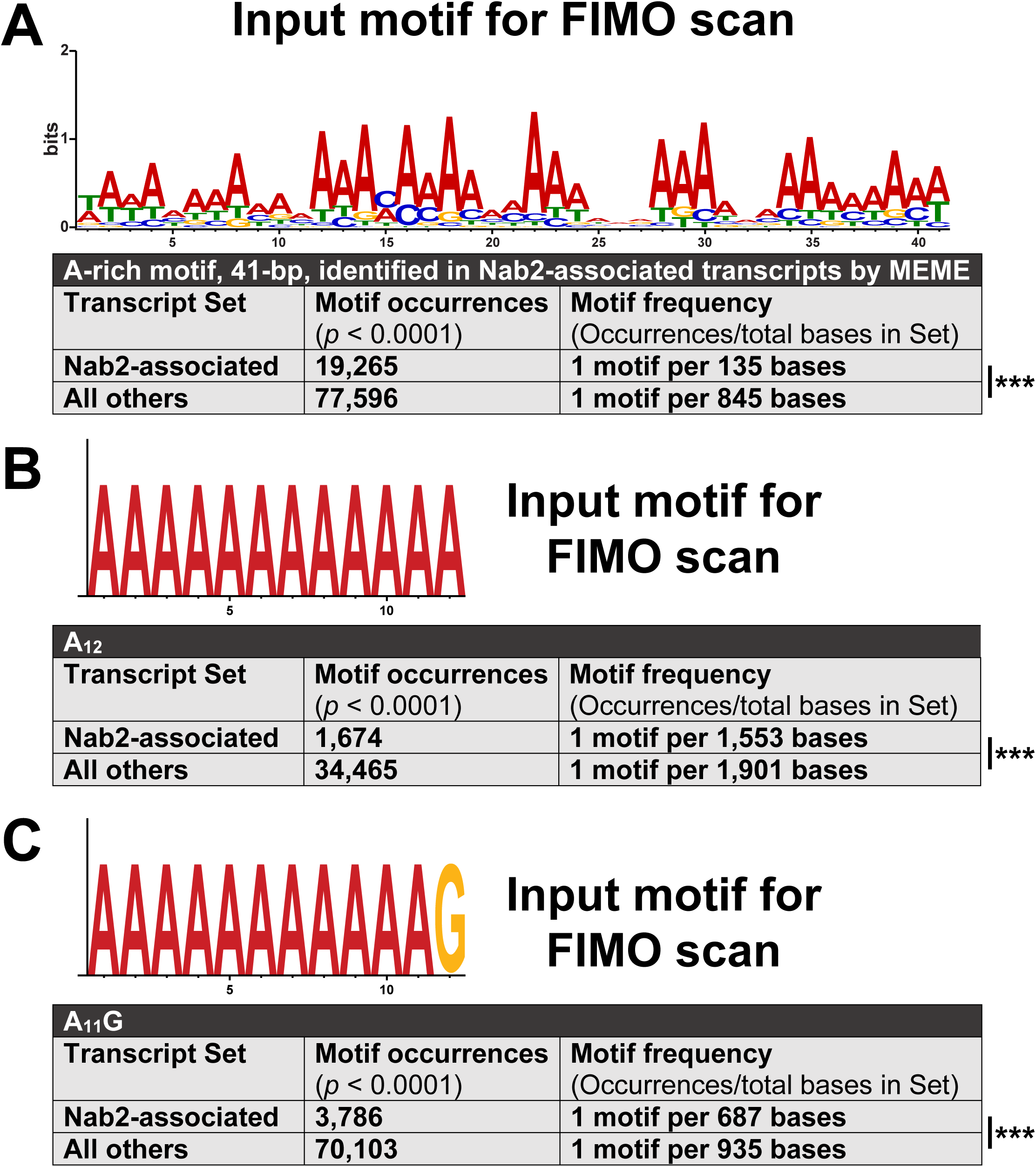
A broad A-rich motif and two specific, canonical Nab2 binding motifs are enriched in Nab2-associated RNAs. Output from transcript set scans by FIMO, which quantifies the occurrences in supplied sequence sets of motifs identical or highly similar to an input motif. Two transcript sets were scanned in each analysis: 1) all transcripts encoded by *Nab2-associated* gene models and 2) all transcripts encoded by *All others*, shorthand for all non-Nab2-associated gene models in the RIP-Seq testable set. (A) A 41-bp A-rich motif, identified by MEME as one of the first ten 6-50 bp motifs within *Nab2-associated* transcripts, was used as input for FIMO. (B) A canonical Nab2 binding motif from *S. cerevisiae*, A_11_G, was used as FIMO input. (C) A simple homopolymer stretch of A’s for which Nab2 would have a very high affinity, A_12_, was used as FIMO input. In all three cases, particularly in (A), the scanned motif is significantly enriched in the *Nab2-associated* transcript set compared to the *All others* transcript set. However, none of the three input motifs are exclusive or nearly exclusive to Nab2-associated transcripts—each is still notably abundant within *All others*. Statistical significance was assessed using the chi-square test (two-sided). ***=*p*<0.001.

As a complement to these analyses, we used FIMO to scan Nab2-associated RNAs for the presence of the smallest canonical binding motifs sufficient for Nab2 association in *S. cerevisiae*—A_12_ and A_11_G (Guisbert *et al*. 2005; Aibara *et al*. 2017). This approach reveals that in *Drosophila* brain neurons A_12_ and A_11_G sites are significantly but moderately more common in Nab2-associated transcripts compared to non-Nab2 associated transcripts. These A_12_ and A_11_G sites appear respectively once every 1,553 and 687 bases on average among Nab2-associated transcripts and once every 1,901 and 935 bases on average among non-Nab2-assoicated transcripts, a 1.2- and 1.4-fold enrichment (Figure 6B,C). Taken together, the findings that Nab2 associates with a specific subset of all RNAs with poly(A) tails, and that these three A-rich motifs are not exclusive to Nab2-associated RNAs, strongly argues that the polyadenosine sequence affinity of Nab2 alone is insufficient to explain Nab2-RNA association specificity in *Drosophila* brain neurons. Other mechanisms must also contribute to Nab2 target choice, such as RNA secondary structure, protein-protein interactions, subnuclear localization, and binding site competition. That said, the significant enrichment of a 41-bp A-rich motif, A_12_, and A_11_G observed in Nab2-associated RNAs suggests Nab2-RNA association is partially mediated through these genetically encoded RNA sequence motifs as well as or instead of through the poly(A) tail.

## Discussion

Mutation of either *ZC3H14 or ATXN2* gives rise to human disease, and the Nab2 and Atx2 RNA-binding proteins encoded by their *Drosophila* orthologs are linked by a shared association with Fmr1 (Sudhakaran *et al*. 2014; Bienkowski *et al*. 2017). Here we show that *Nab2* and *Atx2* interact in multiple contexts in *Drosophila*, specifically in fated eye cells, adult viability, and mushroom body neuronal morphology. Notably, these interactions are dose-sensitive, as heterozygosity for *Atx2* loss-of-function alleles is sufficient to suppress *Nab2* null phenotypes in adult viability and MB morphology. That is, loss of Nab2 may sensitize these domains to reduced Atx2 activity, suggesting these RBPs regulate some common processes. We find that these *Nab2*-*Atx2* interactions are likely not explained by extended, simultaneous co-occupancy of Nab2 and Atx2 in common RNP complexes on shared RNA transcripts. Each protein is concentrated in distinct subcellular compartments in adult mushroom body neurons *in vivo*, and Nab2 and Atx2 weakly associate by co-IP from brain neurons. Thus, to explore an alternative possibility—sequential regulation of shared RNA transcripts—we have carried out the first high-throughput identification of Nab2- and Atx2-associated RNAs in *Drosophila*. We find these proteins associate with overlapping sets of transcripts in *Drosophila* neurons, consistent with their shared and distinct functions and supporting the model of sequential regulation. Identification of these protein-transcript associations promises further insight into the functions shared between and unique to each RBP. In addition, the identification of *Drosophila* Nab2-associated RNAs begins to address longstanding questions about Nab2 function and the particular sensitivity of neurons to Nab2 loss, revealing that Nab2 associates with a specific subset of polyadenylated RNAs *in vivo* despite the theoretical potential to bind across all polyadenylated transcripts suggested by its high polyadenosine affinity *in vitro* (Pak *et al*. 2011).

### A model of opposing regulatory roles for Nab2 and Atx2

We show that Nab2 and Atx2 share associated RNAs in *Drosophila* neurons (Figures 4,5) and that *Atx2* loss-of-function alleles suppress phenotypes of Nab2 loss (Figures 1,2). Taken together, these findings imply that, at least for transcripts crucial for adult survival and MB α lobe morphology, Nab2 and Atx2 exert opposing regulatory roles on their shared associated RNAs. This opposing role possibility aligns with some of the known functions of each protein. Namely, in *S. cerevisiae* Nab2 contributes to proper nuclear processing events including protection from enzymatic degradation, poly(A) tail length control, splicing, and transcriptional termination while also facilitating poly(A) RNA export from the nucleus (Green *et al*. 2002; Hector *et al*. 2002; Kelly *et al*. 2010; Schmid *et al*. 2015; Soucek *et al*. 2016; Alpert *et al*. 2020). If *Drosophila* Nab2 also performs some or all of these nuclear processing roles on its associated RNAs, then Nab2 binding should contribute to transcript stability, nuclear export, and ultimately protein expression. Atx2, in contrast, is a key regulator of translational efficiency in the cytoplasm, suppressing the translation of some target RNAs and activating the translation of others (reviewed in Lee *et al*. 2018). As our data suggest Nab2 and Atx2 act in functional opposition on a shared transcript set, we propose Atx2 primarily functions as a translational inhibitor rather than activator on shared Nab2- and Atx2-associated RNAs. In this model (Figure 7), Nab2 and Atx2 would act in temporal and spatial sequence to balance protein expression from their shared associated RNAs in neurons, with Nab2 promoting proper nuclear RNA processing, stability, and export and Atx2 inhibiting RNA translation, respectively.

**Figure 7.**
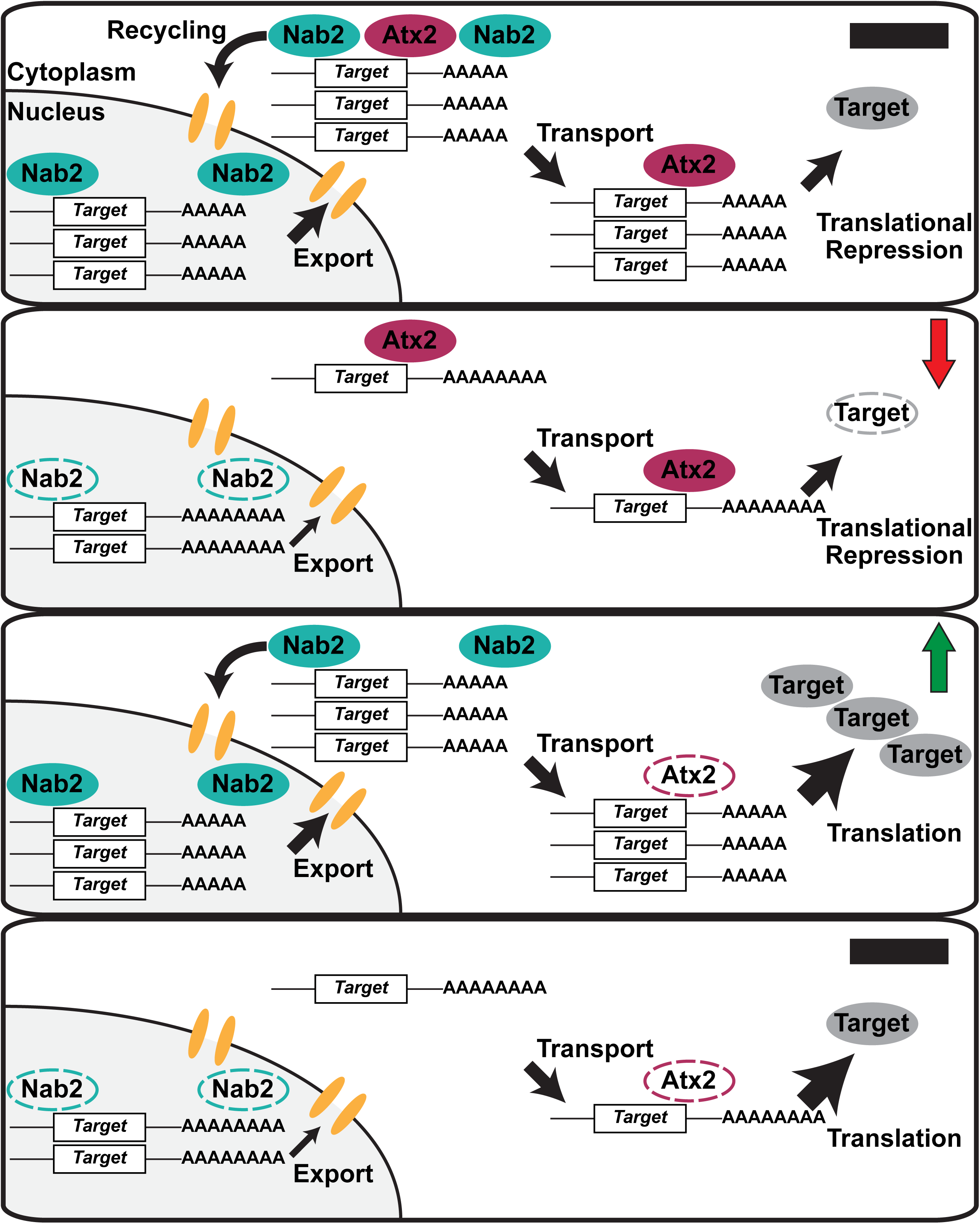
A model of opposing regulatory roles for Nab2 and Atx2 on shared associated RNA transcripts. Nab2 and Atx2 associate with a shared set of RNA transcripts in *Drosophila* brain neurons but primarily localize to separate subcellular compartments and weakly physically associate. *S. cerevisiae* Nab2 regulates nuclear processing, including transcript stability, poly(A) tail length, and export, across a broad RNA transcript set—*Drosophila* Nab2 may perform similar functions on its comparatively limited associated RNA set. Atx2 serves numerous roles in post-transcriptional regulation, including as a miRNA-machinery linked translational repressor. Taken together, these data imply the following model. (*Top*) In the wild-type condition, Nab2 protects transcripts from degradation, limits poly(A) tail length, and contributes to *Target* RNA export from the nucleus, shuttling with its associated transcripts into the cytoplasm. Nab2 and Atx2 may co-occupy the same transcripts briefly or occasionally during nuclear-cytoplasmic mRNP remodeling and prior to Nab2 recycling into the nucleus. Atx2 accompanies *Target* transcripts through transport to their destinations (e.g. synaptic terminals) and contributes to miRNA-mediated translational repression, which is released under certain conditions (e.g. synaptic activity), ultimately contributing to regulated production of wild-type levels of Target protein (black —). (*Second panel*) In *Nab2^ex3^* nulls, *Target* mRNAs are less stable, exhibit longer poly(A) tails, and are exported less efficiently from the nucleus. As a result, less *Target* mRNA reaches its appropriate destination, resulting in a decrease in steady-state levels of Target protein (red ↓). (*Third panel*) In Atx2 loss-of-function heterozygotes (i.e. *Atx2^DG08112/+^* or *Atx2^X1/+^*), less Atx2 protein is expressed and available to repress *Target* translation, resulting in less responsive, higher steady-state levels of Target protein (green ↑). (*Bottom*) Effects of the complete loss of Nab2 in *Nab2^ex3^* and the decrease of functional Atx2 in *Atx2* loss-of-function heterozygotes balance one another. While nuclear *Target* mRNA is less stable and less is exported from the nucleus successfully, these RNAs are also under less strict translational control in partial absence of Atx2, ultimately resulting in Target protein levels and corresponding phenotypes more similar to the wild-type condition (black —). This model represents a prediction from our data and the published knowledge of the functions of each protein—it must be tested in future research, a task enabled by the identification of Nab2- and Atx2-associated transcripts in the current study.

This model of sequential temporal and spatial regulation aligns with evidence that Nab2 and Atx2 primarily localize to different subcellular compartments in adult MBs at steady state and exhibit a low level of co-precipitation from brain neurons (Figure 3). Potential explanations for the combination of distinct localization profiles and limited physical association between Nab2 and Atx2 are found in proposals that *S. cerevisiae* Nab2 shuttles out of the nucleus with bound RNAs during export before releasing them and returning to the nucleus (Aitchison *et al*. 1996; Lee and Aitchison 1999; Duncan *et al*. 2000). Thus, Nab2 and Atx2 may physically share associated RNAs briefly if neuronal *Drosophila* Nab2 similarly shuttles and both RBPs are present during the nuclear-cytoplasmic handoff of mRNP remodeling that follows mRNA export from the nucleus (reviewed in Müller-McNicoll and Neugebauer 2013; Chen and Shyu 2014). Functional and physical links between Nab2 and an RBP with a prominent cytoplasmic localization pattern like Atx2 have been observed previously, specifically with Fmr1 (Bienkowski *et al*. 2017). However, the physical associations observed between Fmr1 and Nab2 are more robust than that observed between Atx2 and Nab2 in the present study (Figure 3B)—this distinction may be partially explained by the different localization patterns of Atx2 and Fmr1. Atx2 is exclusively cytoplasmic in neurons except under certain pathogenic conditions (Huynh *et al*. 2003; Lessing and Bonini 2008; Elden *et al*. 2010), while Fmr1 shuttles between the two compartments, associating with at least some of its target RNAs in the nucleus (Tamanini *et al*. 1999; Kim *et al*. 2009). Thus, Nab2 and Fmr1 may theoretically co-occupy and coregulate shared transcripts in both cellular compartments while Nab2 and Atx2 sequentially regulate shared transcripts exchanged during a nuclear-cytoplasmic handoff, representing two distinct modes of functional interaction between Nab2 and a cytoplasmic RBP.

This model provides a firm foundation and raises many readily testable hypotheses to be explored in future research. The model predicts that for shared Nab2- and Atx2-associated RNAs, loss of Nab2 decreases transcript stability, impedes proper nuclear processing events including poly(A) tail length control, and impairs poly(A) RNA export from the nucleus, ultimately leading to decreases in protein product. Conversely, we predict partial loss of Atx2 releases translational inhibition on these shared transcripts and induces increases in protein product. Finally, loss of both proteins would balance these effects, resulting in steady-state levels of protein product more similar to the wild-type condition. With the identity of Nab2- and Atx2-associated RNAs in hand, future research is enabled to test these predictions.

### Prominent Nab2- and Atx2-associated transcripts provide links to brain development and function

Of all the RBP-associated transcripts identified here, we defined the prominent shared and RBP-specific associated transcripts as those annotated within overrepresented GO terms (Figure 4F, Supplemental Figure 4) and/or passing a 1.5-fold enrichment threshold. The identities and functional roles of these prominent RBP-associated transcripts (examples in Figure 5) provide potential mechanistic explanations for the biological roles of each RBP. For example, the effects of Nab2 and Atx2 on MB morphology may be mediated in part through regulation of shared mRNAs *sm* and *stai*, which respectively encode an hnRNP and a tubulin binding protein both linked to axonal morphology and development (Layalle *et al*. 2005; Lachkar *et al*. 2010; Zwarts *et al*. 2015). The effects of Nab2 and Atx2 on memory (Sudhakaran *et al*. 2014; Kelly *et al*. 2016) may be due in part to regulation of shared transcripts *drk*, *shi*, *Gαo*, and *me31B*, all of which encode proteins with roles in memory formation or retrieval (Dubnau *et al*. 2001; Ferris *et al*. 2006; Moressis *et al*. 2009; Sudhakaran *et al*. 2014). Both Nab2 and Atx2 may be involved in RNAi at multiple levels, regulating *me31B* RNA in neurons in addition to associating, in the case of Atx2, with me31B protein (Lee *et al*. 2017; Bakthavachalu *et al*. 2018). Finally, the suppression of *GMR>Nab2* by *Atx2* alleles in the eye may be explained in part by the shared association of Nab2 and Atx2 with *HmgZ* (*HMG protein Z*) RNA, which encodes a chromatin remodeler linked to survival of R7 retinal photoreceptor neurons (Kanuka *et al*. 2005; Ragab *et al*. 2006).

Among the associated RNAs specific to each RBP, we found only Nab2 associated with *fwe* (*flower*), *Arpc2*, *side-II*, and *SLC22A* RNA, connections which may further explain the role of Nab2 in guiding MB morphology and regulating learning and memory. These transcripts respectively encode a transmembrane mediator of neuronal culling in development (Merino *et al*. 2013), a component of the neuronal growth cone advance-regulating Arp2/3 complex (Hudson and Cooley 2002; Yang *et al*. 2012), an immunoglobulin superfamily member potentially contributing to axon guidance and synapse formation in the optic lobe (Tan *et al*. 2015), and a transmembrane acetylcholine transporter localized to MB dendrites and involved in suppressing memory formation (Gai *et al*. 2016). On the other hand, the association of Atx2 with Atx2-specific RNAs *Xpc* and *tea*, which respectively encode players in the fundamental cellular processes of DNA repair and telomere protection (Henning *et al*. 1994; Goosen 2010; Zhang *et al*. 2016), may partially explain why Atx2 genomic loss, unlike Nab2 genomic loss, is larval lethal (Satterfield *et al*. 2002). In summary, defining the potential functional impact of Nab2- and Atx2-RNA associations like these will provide critical insight into the roles of Nab2 and Atx2 in neurodevelopment and *Drosophila* disease models.

### Nab2 associates with a more specific set of RNAs in metazoans than in *S. cerevisiae*

The degree of RNA association specificity metazoan Nab2/ZC3H14 exhibits has been a longstanding question, in part because competing answers are suggested by the functional similarities and differences between metazoan Nab2/ZC3H14 and the *S. cerevisiae* Nab2 ortholog. In *S. cerevisiae*, Nab2 is essential for viability (Anderson *et al*. 1993) and is a central player in post-transcriptional regulation of many transcripts, serving as a nuclear poly(A)-binding-protein regulating transcript stability (Schmid *et al*. 2015), poly(A) tail length, and poly(A) RNA export from the nucleus among other processes (reviewed in Moore 2005; Chen and Shyu 2014; and Stewart 2019). However, in metazoans Nab2 or the full-length form of ZC3H14 is dispensable for cellular viability, and the effects of either protein on poly(A) tail length and poly(A) RNA export from the nucleus are either less pronounced and likely exerted on fewer transcripts than in *S. cerevisiae* or are not detected (Farny *et al*. 2008; Kelly *et al*. 2010; Wigington *et al*. 2016; Bienkowski *et al*. 2017; Rha *et al*. 2017b; Morris and Corbett 2018). Consistently, Nab2/ZC3H14 have not been found to associate with all polyadenylated RNAs tested in metazoans so far (Wigington *et al*. 2016; Bienkowski *et al*. 2017; Morris and Corbett 2018), but the possibility has remained that these few identified non-Nab2/ZC3H14-associated transcripts are outliers and metazoan Nab2/ZC3H14 associates with a large majority of polyadenylated RNAs similarly to *S. cerevisiae* Nab2 (Tuck and Tollervey 2013), likely in part by binding poly(A) tails. Indeed, the identities of Nab2- or ZC3H14-associated RNAs in metazoans had never previously been addressed with a comprehensive, high-throughput method.

Our results identify a specific set of transcripts that neuronal Nab2 associates with in *Drosophila*. Of the 5,760 transcripts tested in the RIP-Seq, only about 2.5% were found to associate with Nab2 in *Drosophila* neurons (Figure 4), a much smaller percentage of the transcriptome than associates with Nab2 in *S. cerevisiae* (Guisbert *et al*. 2005; Batisse *et al*. 2009; Tuck and Tollervey 2013). Importantly, this likely represents an undercount of all Nab2-associated transcripts in neurons *in vivo*—some RNAs associated with Nab2 in prior studies are absent from our Nab2-associated transcript set (Bienkowski *et al*. 2017; Jalloh *et al*. 2020), and technical limitations impacted our sequencing read depth (see *Methods*). Higher sensitivity approaches (e.g. CLIP-Seq) could reveal a broader set of Nab2-associated transcripts in *Drosophila* than we define here. Nonetheless, in the present study the majority of both RNAs (Figure 4) and tested polyadenosine-rich sequence motifs (Figure 6) were not found to be associated with Nab2, strongly supporting a model in which Nab2 associates with a specific subset of RNAs in *Drosophila* neurons. Perhaps for this more select group of transcripts Nab2 still plays a key role in transcript stability, poly(A) tail length control, transcription termination, and poly(A) RNA export from the nucleus, such that defects will only be observed in targeted examinations of single transcripts and not in bulk assays—one does not always reflect the other (Kelly *et al*. 2014; Bienkowski *et al*. 2017). This model of Nab2 specificity in *Drosophila* aligns well with the knowledge that Nab2/full-length ZC3H14 is essential for cellular viability in *S. cerevisiae* (Anderson *et al*. 1993) but not in *Drosophila* (Bienkowski *et al*. 2017), mice (Rha *et al*. 2017b), or, seemingly, humans (Pak *et al*. 2011; Al-Nabhani *et al*. 2018). This diminished global requirement for Nab2/ZC3H14 in metazoans may be due, at least in part, to functional overlap with PABPN1, an evolutionarily distinct nuclear polyadenosine RNA-binding protein that is absent in *S. cerevisiae* (Mangus *et al*. 2003) but controls poly(A) tail length and is essential in *Drosophila* (Benoit *et al*. 2005), mice (Vest *et al*. 2017), and humans (Hart *et al*. 2015).

The model of Nab2 specificity in *Drosophila* does not conflict with its affinity for polyadenosine, which could theoretically allow Nab2 to bind all transcripts with a poly(A) tail. Even in *S. cerevisiae*, the broad binding profile of Nab2 (Batisse *et al*. 2009) and central role in poly(A) tail length control (Kelly *et al*. 2010), poly(A) RNA export from the nucleus (Green *et al*. 2002), and protection of poly(A) RNA from degradation (Schmid *et al*. 2015) does not translate to binding the poly(A) tails of all transcripts (Guisbert *et al*. 2005; Tuck and Tollervey 2013). More broadly, linear sequence motifs alone are insufficient to explain RBP specificity—RBPs do not generally occupy all of their available binding motifs throughout the transcriptome (Li *et al*. 2010; Taliaferro *et al*. 2016). Moreover, non-paralog RBPs with substantially overlapping or identical linear target motifs still bind distinct RNA target sets, demonstrating that linear motifs are only one of a set of RNA features that direct RBP-RNA associations (Dominguez *et al*. 2018). Based on the present study, these general features of RBPs hold for Nab2 as well. MEME and FIMO motif analyses reveal a long A-rich motif and the canonical Nab2 binding motifs A_12_ and A_11_G are enriched in but not exclusive to Nab2-associated RNAs (Figure 6). Given the behavior of other RBPs, it is consistent that *Drosophila* Nab2 exhibits this binding specificity and, given our RIP-Seq data and previous studies, likely binds some but almost certainly does not bind not all poly(A) tails in *Drosophila* despite its high affinity for polyadenosine RNA *in vitro* (Pak *et al*. 2011).

Taken together, these data align with the model that in metazoans Nab2/ZC3H14 is more specific in its transcript associations than in *S. cerevisiae*. With this model in mind and the Nab2-associated transcripts identified in this study in hand, future research will be enabled to focus on how Nab2 functions on these particular transcripts in *Drosophila*, and why this function is so crucial for adult viability, neuronal morphology, locomotion, and learning and memory. Given that a polyadenosine-rich motif along with A_12_ and A_11_G motifs are correlated with but are not sufficient for Nab2-RNA association, future research must also focus on what additional features of transcripts or their associated proteins promote or prevent Nab2 association.

## Conclusion

In sum, the data we present here identify functional interactions between Nab2 and Atx2 in *Drosophila* brain morphology and adult viability and define a set of RNA transcripts associated with each protein in brain neurons. Crucially, theses RNA sets overlap—some associated transcripts are shared between Nab2 and Atx2 and some are specific to each RBP. Identifying these RBP-associated transcripts provides potential mechanistic links between the roles in neuronal development and function their encoded proteins perform, Nab2, and Atx2. This foundation will be especially important for Nab2, as the exact molecular function of metazoan Nab2/ZC3H14 on the vast majority of its associated RNAs in any cell type remains largely unknown. The identity of many *Drosophila* Nab2-associated transcripts, now revealed, will be required to define Nab2/ZC3H14 function in metazoans and enable our understanding of why loss of this largely nuclear polyadenosine RNA-binding protein results in neurological or neurodevelopmental deficits in flies and mice and in intellectual disability in humans.

## Acknowledgements

The authors would like to thank current and past members of the Moberg and Corbett lab groups, especially Drs. Ayan Banerjee, Rick Bienkowski, Daniel Barron, Binta Jalloh, Stephanie Jones, Annie McPherson-Davie, Milo Fasken, and Sara Leung for their support of, instruction in, and enlightening discussions of this work. We would also like to thank Drs. Bing Yao, Jingjing Yang, Michael Christopher, and Carlos Moreno for initial bioinformatics advice and the Georgia Genomics and Bioinformatics Core (GGBC) at the University of Georgia, especially Tyler James Simmonds and Dr. Magdy S. Alabady, for essential library preparation, sequencing, and assistance in sequencing experiment design and preparation.

We would like to thank Drs. Nancy Bonini and Michael Parisi for the gift of a *Atx2^X1^* stock; Dr. Ravi Allada, Khadijah Hamid, and Dr. Satya Surabhi for the gift of a *UAS>Atx2-3xFLAG* stock; Dr. Chunghun Lim and for the gift of rabbit α-Atx2; Dr. Corey S. Goodman for the contribution of 1D4 Anti-Fas2 to the DSHB; Drs. Gary Bassell, Roger Deal, Steven Warren, James Q. Zheng, and their respective labs for assistance and use of their equipment; Laura Fox-Goharioon for confocal microscope training; Dr. Michael I. Love for extensive public online instruction in the methodology and use of *DESeq2*; Dr. Mauricio Rodriguez-Lanetty for a public TRIzol-column hybrid RNA extraction protocol; and Eileen Chow for public video instruction in bulk *Drosophila* head isolation.

The authors would also like to thank with particular enthusiasm the authors, contributors, and ongoing maintainers of the incredible public resources supporting this work and without which it would not have been possible, including Flybase (NIH U41HG000739, UK MRC MR/N030117/1), the Galaxy Project (NIH 2U41HG006620), the R Project, the Developmental Studies Hybridoma Bank (University of Iowa, NIH), and the Bloomington Drosophila Stock Center (NIH P40OD018537). This research was funded by grants from the National Institutes of Health, specifically from the National Institute of Child Health and Human Development (F31 HD088043) to J.C.R. and from the National Institute of Mental Health (R01 MH107305) to A.H.C. and K.H.M. The authors declare no conflicts of interest.

